# Minimization of the E. coli ribosome, aided and optimized by community science

**DOI:** 10.1101/2023.11.09.566451

**Authors:** Tiyaporn Tangpradabkul, Michael Palo, Jill Townley, Kenneth B. Hsu, Eterna participants, Sarah Smaga, Rhiju Das, Alanna Schepartz

## Abstract

The ribosome is a ribonucleoprotein complex found in all domains of life. Its role is to catalyze protein synthesis, the messenger RNA (mRNA)-templated formation of amide bonds between α-amino acid monomers. Amide bond formation occurs within a highly conserved region of the large ribosomal subunit known as the peptidyl transferase center (PTC). Here we describe the stepwise design and characterization of mini-PTC 1.1, a 284-nucleotide RNA that recapitulates many essential features of the *Escherichia coli* PTC. Mini-PTC 1.1 folds into a PTC-like structure under physiological conditions, even in the absence of r-proteins, and engages small molecule analogs of A- and P-site tRNAs. The sequence of mini-PTC 1.1 differs from the wild type *E. coli* ribosome at 12 nucleotides that were installed by a cohort of citizen scientists using the on-line video game Eterna. These base changes improve both the secondary structure and tertiary folding of mini-PTC 1.1 as well as its ability to bind small molecule substrate analogs. Here, the combined input from Eterna citizen-scientists and RNA structural analysis provides a robust workflow for the design of a minimal PTC that recapitulates many features of an intact ribosome.

## INTRODUCTION

The ribosome is a ∼2.3-4.3 MDa ribonucleoprotein complex that catalyzes messenger RNA (mRNA)-templated amide bond formation during protein synthesis. It is conserved across all domains of life. In bacteria such as *E. coli*, the ribosome is composed of three ribosomal RNA (rRNA) molecules and more than fifty ribosomal proteins (r-proteins), and its 2.8 MDa structure has been characterized in extraordinary detail (1) (**Figure 1A**). These structures reveal a 30S small ribosomal subunit and a 50S large ribosomal subunit that each play a distinct role in protein synthesis. The small ribosomal subunit, which contains the 16S rRNA and 21 r-proteins, decodes the mRNA to define protein primary sequence, whereas the large subunit, which contains the 23S rRNA, 5S rRNA, and 33 r-proteins, catalyzes peptide bond formation. Peptide bond formation occurs within a highly conserved region of the 23S rRNA known as the peptidyl transferase center (PTC) (**Figure 1B**). The PTC is considered the most ancient region of the modern ribosome and is believed to predate the last universal common ancestor (LUCA) (2, 3). It is now well appreciated that the ribosome is a ribozyme, as the RNA components of the PTC, rather than the r-proteins, promote nucleophilic attack of the α-amino group of the aminoacyl-tRNA on the ester carbonyl of the peptidyl-tRNA (4–6). In this work, we sought to establish whether an isolated PTC can fold into a discrete ribosome-like structure, and whether this folding could be improved *via* game play by citizen scientists.

**Figure 1.**
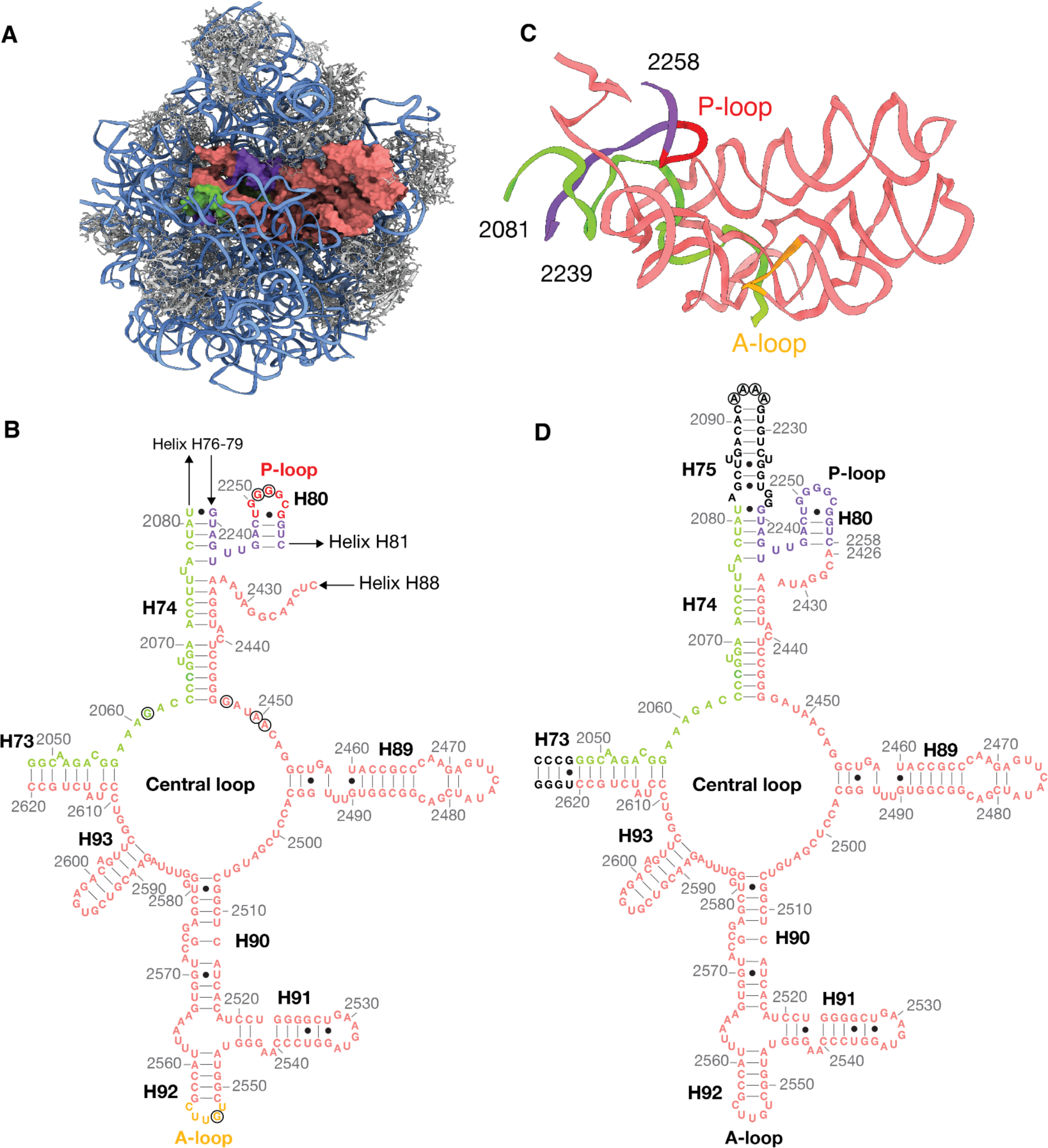
Design of a first generation mini-PTC (WT mini-PTC). (**A**) Cartoon illustrating the cryo-EM structure of the *E.coli* 50S ribosomal subunit (PDB: 7K00) visualized using Chimera X 1.5, viewed from the subunit interface. The 23S rRNA and 5S rRNA are shown as blue ribbons, and ribosomal proteins are shown in gray. The PTC is shown in green, purple, and pink to highlight three noncontiguous segments of the PTC, respectively. (**B**) Secondary structure of domain V of the *E. coli* 23S rRNA, showing the central loop and nearby helices and stems. Residue numbers and helices are provided in gray and black, respectively. Bases shown in green, purple, and pink correspond to the PTC region shown in (**C)**. P-loop and A-loop shown in (**C)** are highlighted in red and yellow, respectively. Highly conserved PTC residues mentioned in the text are highlighted within the black open circles. (**C**) Ribbon structure of the PTC viewed from the same orientation as in (**A**). (**D**) Secondary structure diagram of the first-generation mini-PTC design. Additional *E. coli* 23S RNA residues that were extended to the construct are shown in black. Arbitrary A4 tetraloop is highlighted within the black open circles. Residue numbers and helices are provided in gray and black, respectively.

In *E. coli,* the PTC assembles *via* the convergence of three non-consecutive regions of the 23S rRNA within domain V. These regions include residues 2048 - 2081 (H73 and H74, shown in green), 2239 - 2264 (H80, shown in purple) and 2422 - 2620 (H89-H93, shown in pink) (**Figure 1B**). When visualized in two dimensions, the PTC consists of a large central loop interrupted by five short helices: H73, H74, H89, H90, and H93 (7). When visualized in three dimensions within a fully assembled ribosome (PDB: 7K00) (1), the PTC consists of a well-defined hairpin with elements of two-fold pseudo-symmetry (**Figure 1C**). Residues on one side of the two-fold axis are associated with the ribosomal P-site, whereas those on the other side are associated with the A-site (8). Inside the semi-symmetrical interface, residues G2061, G2447, A2450, and A2451 form a network of hydrogen bonds to position the P-site peptidyl-tRNA and the A-site aminoacyl-tRNA for amide bond formation (4). tRNA substrates are further stabilized by base-pairing interactions with neighboring PTC residues. These include G2251 and G2252 of the 23S rRNA P-loop (2249-UGGGGCG-2255, circled and shown in red), which base-pair with C75 and C74 of the P-site tRNA, as well as G2553 of the 23S rRNA A-loop (2552-UGUUC-2556, circled and shown in yellow), which base pairs with C75 of the A-site tRNA (**Figure 1B, C**) (9, 10). The structurally complex interactions within the PTC are essential for ribosome activity and are highly conserved across all kingdoms of life (3, 11, 12). Indeed, efforts to alter residues within the PTC to favor new bond-forming reactions have met with limited success (13–15).

Previous reports have identified small PTC-derived RNA subdomains that exhibit low levels of peptidyl transferase activity. Wang and coworkers reported mass spectrometry results that imply the production of an ennea-lysine oligomer upon incubation of lysine-charged tRNA fragments with a dimer of a 108-nucleotide (nt) PTC-derived rRNA subdomain from *T. thermophilus* (16, 17). Yonath and coworkers reported several 67-135-nt PTC-derived rRNA subdomains derived from *T. thermophilus*, *S. aureus*, and *E. faecium* that act as dimers to promote peptide bond formation between small molecule P-site and A-site substrate analogues (18). In addition, Tamura and coworkers reported 70 and 74-nt PTC-derived rRNA subdomains from *D. radiodurans* that bind to alanyl-tRNA minihelices and promote the formation of an Ala-Ala dimer (19). These previous studies are important because they imply that the proto-ribosome evolved via the dimerization of a small PTC-derived rRNA subdomain. However, because the small rRNA subdomains were not structurally characterized, before or while bound to ribosomal substrates, they shed no light on the extent to which the rRNA subdomains recapitulate the fold of the modern-day PTC.

Here we combine rational design and community science game play to identify a small rRNA subdomain that assembles as a single unit into a secondary and tertiary fold that mimics that of the *E. coli* PTC. A first-generation design developed *de novo* was subsequently improved through a collaboration with individuals who play Eterna. Eterna is an internet-scale citizen science game that challenges players to design RNA sequences that fold into predetermined secondary structures (20). During the game, players design sequences by exploring changes in Watson-Crick base-pairing, which are subsequently evaluated computationally and via direct biochemical analysis. This crowd-sourcing approach to discovery leverages both human insights and a massive number of unique strategies to solve a complex RNA folding problem. Eterna player-guided RNAs can outperform computational algorithms and generate unique RNA designs, such as mRNAs with increased stability or efficient and reversible riboswitches (21, 22). Recent work has used Eterna as a platform to generate ribosomes with 23S and 16S rRNA mutants that increase translation activity relative to the WT sequence when evaluated *in vitro* (23).

Here we describe the stepwise design and characterization of mini-PTC 1.1, a 284-nucleotide RNA that recapitulates many essential features of the *E. coli* ribosomal PTC. Mini-PTC 1.1 folds into a PTC-like structure under physiological conditions, even in the absence of r-proteins and assembly factors and engages small molecule analogs of A- and P-site tRNAs. The sequence of mini-PTC 1.1 differs from the wild type *E. coli* ribosome at 12 bases, all of which were identified by an Eterna player. These base changes, located throughout domain V, improve both secondary structure and tertiary folding and resulted in substantially improved binding to small molecule analogs of A- and P-site tRNAs. The combined Eterna citizen-scientist/RNA structural analysis platform allows for robust design of a minimal PTC construct with desired structure.

## MATERIALS AND METHODS

### RNA synthesis, purification, and folding

For each mini-PTC variant, two types of RNA constructs were synthesized for use in different downstream experiments. 284-nt mini-PTC constructs were prepared for native gel electrophoresis experiments, whereas 357-nt mini-PTC constructs containing reference hairpins at the 5′ and 3′ ends of the mini-PTC sequence were prepared for SHAPE analysis. For 284-nt mini-PTC RNAs that have the sequence starting with a U at 5′ end, including mini-PTC 1.1, 1.2, and 1.7, the cis-acting 5′ hammerhead ribozyme has been extended at the 5′ ends of the mini-PTC sequence to optimize the yield for *in vitro* transcription (referred to as 284-nt mini-PTC-transzyme constructs) (24). DNA and RNA sequences used to generate 357-nt mini-PTC, 284-nt mini-PTC, and 284-nt mini-PTC-transzyme constructs are shown in Supplementary Data.

All DNA primers were purchased from Integrated DNA Technologies. Primers for PCR assembly of DNA transcription templates were designed using the Primerize web server (25). PCR assembly was performed using the Q5 High-Fidelity 2X Master Mix (New England Biolabs), 4 µM of 5′ terminal primer, 4 µM of 3′ terminal primer, and 0.04 µM of internal primers. The thermocycler setting consisted of 30 cycles of 98 °C for 10s, 60 °C for 30s, 72 °C for 30s. PCR products were then purified using QIAquick PCR Purification Kit (Qiagen) according to the manufacturer’s protocol and quantified concentration using NanoDrop ND-1000 device (Thermo Scientific). 357-nt mini-PTC and 284-nt mini-PTC RNA constructs were prepared by *in vitro* transcription using the TranscriptAid T7 High Yield Transcription Kit (Thermo Scientific) according to the manufacturer’s protocol. 284-nt mini-PTC-transzyme constructs were prepared by *in vitro* transcription using a modified version of a published procedure (26). Transcription reactions (200 µl) contained the following components: 50 mM Tris-HCl (pH 8.1), 0.01% Triton X-100, 100 mM DTT, 2 mM spermidine, 10 mM ATP, 10 mM CTP, 10 mM GTP, 10 mM UTP, 50 mM MgCl_2_, 10 ng µl^−1^ DNA template, 1 unit ml^−1^ *E. coli* inorganic pyrophosphatase (NEB), and 0.2 mg ml^−1^ in-house T7 RNA polymerase. The reaction mixtures were incubated at 37 °C in a thermocycler for 16 h and subjected to additional incubation at 60 °C for 2 h to cleave off the desired mini-PTC RNA.

After *in vitro* transcription, 357-nt mini-PTC RNA constructs were purified with RNA Clean and Concentrator-25 columns (Zymo Research) according to the manufacturer’s protocol, while 284-nt mini-PTC RNA and 284-nt mini-PTC-transzyme constructs analyzed by native gel electrophoresis were additionally purified on a 5% 19:1 acrylamide:bis, 7 M urea polyacrylamide gel. Then, a 2× stop solution (80% formamide, 50 mM EDTA, 0.025% xylene cyanol, and 0.025% bromophenol blue) was added to a final concentration of 1× and the samples were loaded on the gel. The gel was run at 150 V for 1 h 30 min, then visualized briefly with a 254-nm UV lamp, held far from the gel to minimize RNA damage (27). The RNA band was excised from the gel and eluted with 1 mL 0.3 M NaOAc pH 5.2 at 4 °C for 16 h. Gel traces were removed by passing the solution through a 0.22 µm syringe filter. The recovered RNA solution was then precipitated with 3 volumes of 100% (v/v) ethanol at −80 °C for at least 30 min. RNA pellet was collected by centrifugation at 21,130 × g for 30 min at 4 °C, washed once with 70% (v/v) ethanol, and resuspended in RNase-free water. RNA was heated to 90 °C for 3 min in 50 mM Na-HEPES pH 8.0 and cooled to room temperature for 12 min. Then, MgCl_2_ was added to a final concentration of 10 mM and incubated at 50 °C for 20 min to refold RNA.

### Native gel electrophoresis

Native gel electrophoresis was performed as described previously (28). Briefly, a 15 mL solution containing 8% 19:1 acrylamide:bis, 10 mM MgCl_2_, 67 mM Na-HEPES pH 7.2, and 33 mM Tris, pH 7.2 was mixed with 150 μL 10% (w/v) ammonium persulfate and 30 μL TEMED and cast in a BioRad Mini-Protean Tetra handcast systems. The gel was pre-run in the native gel running buffer (10 mM MgCl_2_, 67 mM Na HEPES pH 7.2, and 33 mM Tris, pH 7.2) for 1 h in a 4 °C cold room. Samples were prepared by incubating 200 ng RNA in 50 mM Na-HEPES pH 8 at 90 °C for 3 min to denature the RNA. The RNAs were cooled to room temperature for 10 min, then MgCl_2_ was added to a final concentration of 10 mM in 8 μL in the folding condition. For denaturing conditions, urea was added to a final concentration of 6 M in 8 μL instead. RNA samples were refolded at room temperature, 37 °C, 50 °C, or 65 °C for 20 min and then cooled to room temperature for 10 min. Each sample was mixed with 2 μL 5× native gel loading buffer (250 mM Na-HEPES pH 7.5, 5 mM EDTA pH 8, 50% glycerol, 0.05% each bromophenol blue and xylene cyanol) and loaded onto the gel. The gel was run at 200 V for 1 h 30 min at 4°C, stained with SYBR Gold dye for 5 min, and destained with the native gel running buffer for 1 min. The gel was visualized on a BioRad ChemiDoc MP Imaging System.

### Chemical probing by selective 2′ hydroxyl acylation analyzed by primer extension (SHAPE)

SHAPE chemical probing was performed with RNA constructs containing reference hairpins as described previously (29, 30). For each mini-PTC construct, SHAPE experiments were done with two no SHAPE modification controls (for background measurements) and three SHAPE modification reactions. SHAPE modification reaction mixtures consisted of 15 µL of 1.2 pmol of RNA in 50 mM Na-HEPES pH 8.0 with 10 mM MgCl_2_ and 5 µL of 19 mM freshly prepared 1-methyl-7-nitroisatoic anhydride (1M7) in DMSO. In control reactions, 5 µL of 50 mM Na-HEPES pH 8.0 with 10 mM MgCl_2_ was added instead of 1M7. The reaction mixtures were incubated at room temperature for 15 min. To quench reactions, 9.8 µL of the premixed solution consisted of 5 µL of 0.5 M Na-MES (pH 6.0), 3 µL of 5 M NaCl, and 1.8 µL water was added to the 20 µL reaction mixtures. RNAs were then recovered by ethanol precipitation. For each 20 µL reaction, the solution containing 20.2 µL water, 0.5 μl 20 µg/ml glycogen, 1 μl 100 mM EDTA pH 8, and 175 μl absolute ethanol was added to precipitate RNA at −80°C for 30 min, followed by centrifugation at 21,130 × g for 30 min at 4°C (31). RNA pellet was resuspended in 4.25 μl of 0.5× TE buffer.

### Primer extension analysis

Primer extension was done by adding 0.5 µL of 2 µM 5′ 6-FAM-labeled primer (AAAAAAAAAAAAAAAAAAAAGTTGTTGTTGTTGTTTCTTT) complementary to the 3′ end of the RNAs. Primer was annealed to the RNAs by incubating at 65 °C for 5 min and then at 4°C for 5 min, then at room temperature for 5 min. The mixtures of modified RNAs and primers were then reverse transcribed by the addition of a reaction mix consisting of 0.25 µL of SuperScript III reverse transcriptase (Invitrogen) 2.5 µL of 5× SuperScript First Strand buffer (Invitrogen), 0.5 µL of 0.1 M DTT (Invitrogen), 2 µL RNase-free water (Sigma-Aldrich), and 2.5 µL of dNTP mix (10 mM each, Invitrogen). For sequencing reactions, the reactions were set up using 0.25 µL of SuperScript III reverse transcriptase (Invitrogen) 2.5 µL of 5× SuperScript First Strand buffer (Invitrogen), 0.5 µL of 0.1 M DTT (Invitrogen), 0.25 µL of 10 mM ddATP, ddTTP, ddCTP, or ddGTP (Trilink Biotechnologies), 3.25 µL RNase-free water (Sigma-Aldrich), and 1 µL of dNTP mix (10 mM each, Invitrogen). The reverse transcription reaction was incubated at 50 °C for 1 h. RNA was then degraded by the addition of 1.5 µL 1 N NaOH (Fisher Scientific) and incubation at 70 °C for 15 min. The solution mixture containing cDNA products was neutralized with 1.5 µL 1 N HCl (Fisher Scientific). To separate cDNA products by capillary electrophoresis, 0.5 µL of the neutralized reaction mixtures were added to 9 µL HiDi Formamide (Applied Biosystems) and 0.5 µL ROX 350 size standard on the 3730XL DNA Analyzers (UC Berkeley DNA sequencing facility).

The electropherogram was analyzed using the HiTRACE software, in which the traces were aligned based on the ROX-350 internal standard coloaded with all samples. Band assignments were done by comparing each band to the sequencing ladders. SHAPE reactivities were calculated by subtracting the reactivities from no modification controls and further normalized with the reactivity of reference hairpins (GAGUA) at the 5′ and 3′ ends (30). SHAPE reactivities then were plotted against the secondary structure diagram using the RiboDraw (https://github.com/ribokit/RiboDraw) (32) and RiboPaint packages (https://ribokit.github.io/RiboPaint/). MATLAB and python scripts used to generate SHAPE analysis and heatmap are available at https://github.com/gem-net/miniPTC-paper.

### SHAPER predictions

The 3D coordinates of the predicted tertiary structure of Mini-PTC WT in PDB format were derived from the *E.coli* fully assembled ribosome (PDB 7K00). The SHAPE experimental data in a .txt file was submitted to the SHAPER web server http://rna.physics.missouri.edu/shaper/ (33). Predicted SHAPE profile was downloaded directly from a web browser. The input structure and the output SHAPE profile data are available in Supplementary Data.

### Distribution identification and outlier analysis

Differential SHAPE reactivity data was fitted to the Cauchy probability density function using the Fitter 1.6.0 python package (https://zenodo.org/record/8226571). To identify the extreme outliers in the distribution, we applied the interquartile range (IQR) method that can be used to measure variability of the data based on quantiles.

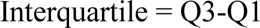

where *Q1* = the first quartile (25% percentile) and Q3 = the third quartile (75% percentile)

Extreme outliers were defined as values that are greater or lower than 3 × IQR Python scripts used to identify the distribution and outliers are available at https://github.com/gem-net/miniPTC-paper.

### SHAPE footprinting in the presence of CCA-pcb and C-pmn

Mixtures containing 1.2 pmol WT mini-PTC or mini-PTC 1.1 RNA, 100 µM CCA-pcb (Horizon Discovery), and 100 µM C-pmn (Horizon Discovery) were heated to 90 °C for 3 min in 50 mM Na-HEPES pH 8.0 and cooled on ice for 5 min. Then, MgCl_2_ was added to a final concentration of 10 mM and incubated at 50 °C for 20 min to fold RNA. SHAPE and primer extension analysis were performed as mentioned above.

### Fragment reaction and RNase A digestion

Fragment reactions were set up using a modified version of a published procedure (18). Reaction mixtures (10 µL) in the presence of mini-PTC 1.1 contained the following components: 20 µM Mini-PTC 1.1, 100 µM CCA-pcb (Horizon Discovery), 100 µM C-pmn (Horizon Discovery), 50 mM HEPES-KOH pH 7.5, 20 mM Mg(OAc)_2_, and 400 mM KCl. To re-fold RNA, 100 µM mini-PTC 1.1 in RNase-free water was heated at 90 °C for 3 min and snap-cooled on ice for 5 min.

RNA was then diluted to 20 µM and mixed with remaining components. The reaction mixtures were incubated at either −20 °C, 4 °C, or 50 °C for 2 or 24 h. 0.1 µM *E. coli* 50S ribosomal subunit (kindly provided by Kristina Boyko) and RNase-free water were used to replace 20 µM mini-PTC 1.1 in the reaction with *E. coli* 50S ribosomal subunit and the control reaction, respectively. Re-folding step was not necessary to perform in the reaction with *E. coli* 50S ribosomal subunit and the control reaction.

Following the reactions, reaction mixtures (10 µL) were digested with 1 µL of 1.5 U/µL RNase A (Millipore-Sigma) and incubated at room temperature for 5 min. Proteins were then precipitated upon addition of 50% trichloroacetic acid (TCA, Sigma-Aldrich) to a final concentration of 5%. Samples were frozen on dry ice for 5 min to precipitate protein, insoluble material was removed by centrifugation at 21,300 × g for 10 min at 4 °C. The soluble fraction was then transferred to autosampler vials. Prior to injection, 1 µL of 40 ng/mL Leu-enkephalin (Waters) was added to each 10 µL reaction as an internal standard. Samples were kept on ice until immediately before LC-MS analysis and returned to ice immediately afterwards.

### LC-MS Analysis

Samples analyzed by mass spectrometry were resolved using a Zorbax Eclipse XDB-C18 RRHD column (2.1 × 50 mm, 1.8 μm, room temperature, Agilent Technologies part # 981757-902) fitted with a guard column (Zorbax Eclipse XDB-C18, 2.1 x 5 mm 1.8 μm, Agilent part # 821725-903) using a 1290 Infinity II UHPLC (G7120AR, Agilent). The mobile phases used were (A) 0.1% formic acid in water; and (B) 0.1% formic acid in 100% acetonitrile. This 11.5-min method used a flow rate of 0.7 mL/min and began with Mobile Phase B held at 5% for 2 min, followed by a linear gradient from 5 to 95% B over 7.5 min, then finally B held at 95% for 2 min. The following parameters were used: fragmentor voltage of 175 V, gas temperature of 300°C, gas flow of 12 L/min, sheath gas temperature of 350°C, sheath gas flow of 12 L/min, nebulizer pressure of 35 psi, skimmer voltage of 65 V, Vcap of 3500 V, and collection rate of 3 spectra/s. Expected exact masses of puromycin-phenylalanine-caproic acid-biotin (m/z: 958.4604) and leucine enkephalin (m/z: 556.2771) were calculated using ChemDraw 19.0 and extracted from the total ion chromatograms ± 100 ppm by LC-HRMS with an Agilent 6530 Q-TOF AJS-ESI (G6530BAR).

The yield of C-pmn-pcb in the fragment reactions was determined by calculating it using the equation below:

Yield = (Peak area under curve of pmn-pcb)/(Peak area under curve of Leu-enk) × 10^3^

## RESULTS

### Design of WT mini-PTC, a first-generation minimal PTC

To develop an RNA-only minimal PTC, we first designed a single RNA strand (**Figure 1D**) containing all domain V nucleotides that interact with one another within the core PTC hairpin (**Figure 1B**). Extraneous segments with other functions were excluded. The three noncontiguous 23S rRNA segments that comprise the PTC core (colored green, purple, and pink in **Figure 1D**) were converted into a single RNA strand upon addition of one stem-loop and one loop-loop ligation. This conversion began by capping H75, which connects the PTC to the L1 stalk comprising helices H76-H79 (34–36), with a stable A4 tetraloop (A2092-A2095) (37). Next, nucleotides at the 3′ end of H80 near the P-loop (2254-CGGUC-2258) were connected directly to the loop 2426-ACGGA-2430 excluding helices H81-H88 that are located in the central protuberance. Finally, four GC-rich base pairs present in the beginning of H73 were included to form the 5′ and 3′ termini of the PTC core. The resulting first-generation mini-PTC (referred to henceforth as WT mini-PTC) contains 284 nucleotides that comprise the central loop, P-loop, and A-loop. Our initial experiments focused on whether this 284-nucleotide, native-sequence RNA contained sufficient information to direct formation of a stable PTC-like secondary structure without need for ribosomal proteins, assembly factors, or RNA chaperones.

### WT mini-PTC moderately recapitulates the native PTC fold

When prepared by T7-catalyzed transcription *in vitro* and purified, WT mini-PTC migrated as a single band on a denaturing acrylamide gel (**Figure S1A**). Two bands were observed on a native acrylamide gel when WT mini-PTC was incubated in 6 M urea and then re-folded at temperatures between 25 and 50 °C (**Figure S1B**). These two bands were also observed on a native gel when WT mini-PTC was heated to 90 °C, cooled down at RT, and re-folded at 25 or 37 °C in the presence of 10 mM MgCl_2_. However, only a single band was observed when WT mini-PTC was heated and re-folded at 50 °C in the presence of 10 mM MgCl_2_ **(Figure S1B**).

These results imply that WT mini-PTC can assemble into at least two structures, but assumes what appears to be a homogenous conformation when refolded at 50 °C in the presence of 10 mM MgCl_2_. For this reason, WT mini-PTC samples used for structural analysis as described below were refolded at 50 °C in the presence of 10 mM MgCl_2_.

Next, we used Selective 2′ Hydroxyl Acylation analyzed by Primer Extension (SHAPE) analysis to define the secondary structure of WT mini-PTC at single-nucleotide resolution (38–41). SHAPE probes the reactivity of ribose 2′ hydroxyl groups towards acylating agents such as 1-methyl-7-nitroisatoic anhydride (1M7) (**Figure 2A)**. Acylation results in covalent adducts that inhibit the activity of reverse transcriptase, leading to truncated DNA products when the modified RNA is reverse-transcribed. The truncated products are then identified and quantified by capillary electrophoresis. In general, high SHAPE reactivity is observed at nucleotides located in loops and other unpaired regions, whereas low SHAPE reactivity is observed at nucleotides within helices and constrained loops (31, 38, 42). To create a construct suitable for quantitative SHAPE mapping, we extended the 5′ and 3′ ends of WT mini-PTC with reference hairpins (GAGUA) to allow data normalization (29, 30) (colored red in **Figure S2A**). To perform a SHAPE experiment, WT mini-PTC was folded at 50 °C in the presence of 10 mM MgCl_2_, cooled to room temperature for 10 min, reacted with 4.7 mM 1M7 for 15 min, and primer-extended into cDNA products using SuperScript III reverse transcriptase. Reaction products were resolved and quantified using capillary electrophoresis (**Figure S2B**). Electropherograms were processed using HiTRACE software (43).

**Figure 2.**
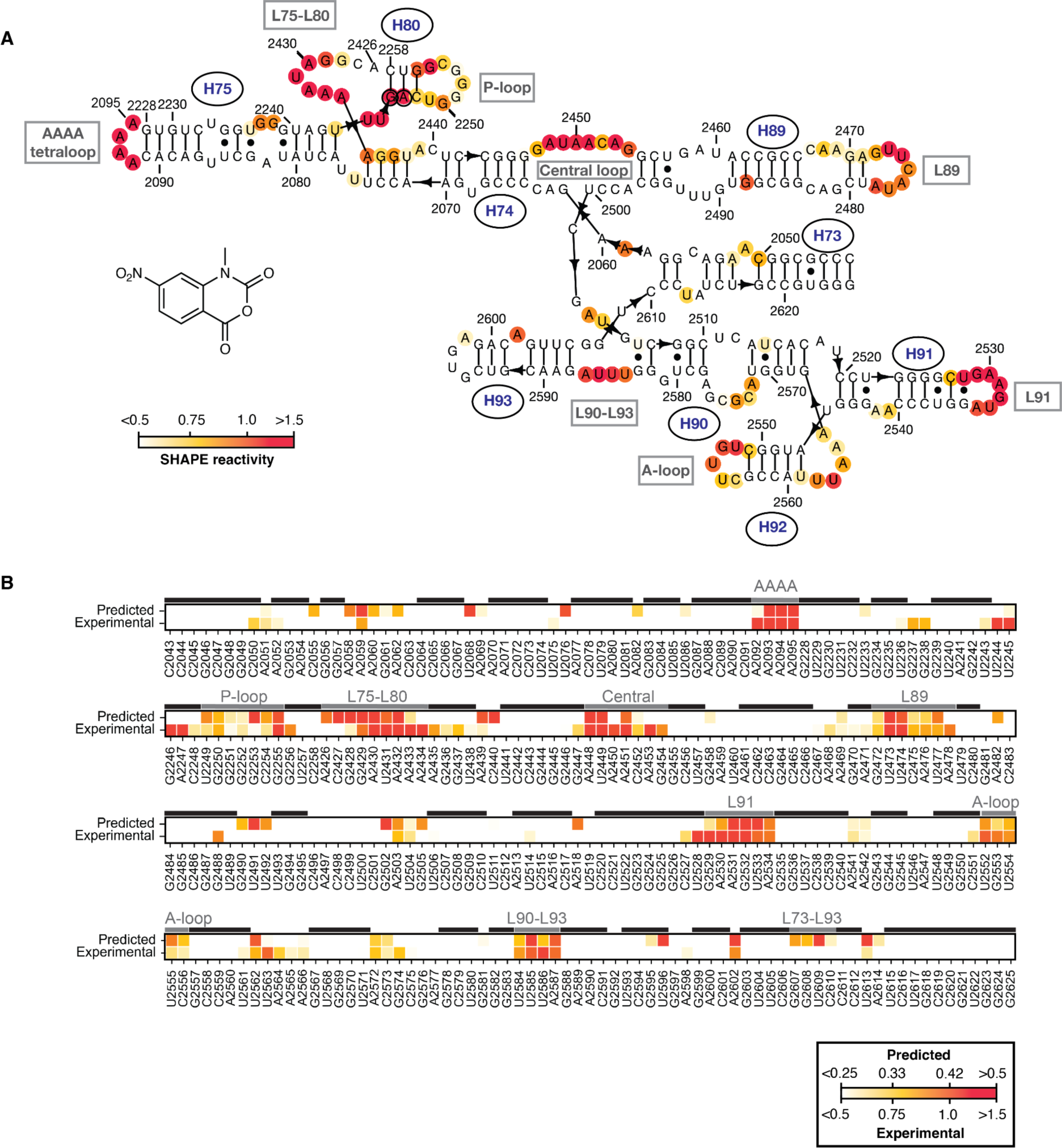
WT mini-PTC recapitulates several structural elements of the ribosomal PTC. **(A)** Structure of 1M7 alongside the SHAPE reactivity of WT mini-PTC plotted as a heatmap on the predicted secondary structure of the *E. coli* 50S ribosomal subunit. Nucleotide positions were indicated using conventional *E. coli* 23S rRNA numbering. **(B)** Heatmap showing the difference between predicted and experimental SHAPE reactivity. SHAPE reactivity was plotted on a red-yellow-white spectrum where red represents high SHAPE reactivity (>0.5 predicted; >1.5 experimental) and white represents low SHAPE reactivity (<0.25 predicted; <0.5 experimental). Variations between experimental and predicted SHAPE reactivity were attributed to higher background in the experimental data. Base-pairing residues and loop regions described in Results were marked with a black line and a gray line, respectively, over the heatmap.

Those nucleotides within WT mini-PTC exhibiting significant 1M7 reactivity over background were mapped on the anticipated PTC secondary structure, as defined by the cryo-EM structure of the *E. coli* 50S ribosomal subunit at 2Å resolution (1), to identify regions of agreement and contradiction **(Figure 2A**). Both the engineered AAAA tetraloop at the terminus of H75 (A2092-A2095) and the homologation between H75 and H80 (A2426-A2434), which is referred to as L75-L80, show high reactivity in most residues (high reactivity is defined as a SHAPE reactivity > 1.5; these residues are colored red in **Figure 2A**). High reactivity is also observed in residues located in L89 (G2472-A2478), L91 (G2529-A2534), L90-L93 (U2584-A2587), the central loop (G2447-2454), P-loop (U2249-G2255), and the A-loop (U2552-C2556). In contrast, little or no reactivity is observed in most Watson-Crick base-paired regions: of the 140 nucleotides that are expected to be Watson-Crick base-paired, only 17 (12.1%) react with 1M7 at a level significantly over background. Only two ostensibly base-paired residues (1.4%), G2246 and A2247 (black circled residues in **Figure 2A**) show high reactivity with 1M7.

To further interrogate the extent to which WT mini-PTC recapitulated the expected PTC secondary and tertiary structure, we compared the noise-adjusted experimental SHAPE reactivity to that predicted by a 3D Structure-SHAPE predictoR (SHAPER), which estimates SHAPE reactivity of a user-defined RNA tertiary structure (**Figure 2B**). SHAPER predicts SHAPE reactivity on the basis of base-pairing, solvent accessibility, nucleotide stacking, and ribose sugar conformations as well as RNA sequence-dependent bias (33). When comparing the SHAPER-predicted reactivity of a 461-nt domain V of 23S RNA to the experimental reactivity previously obtained from native *E. coli* 23S RNA (44), a correlation of 0.58 was observed between the two datasets (**Figure S3**). Previous work has shown that SHAPER can be used to predict the native structure of a 29-nt HIV TAR RNA (45) with a correlation score of 0.88, while near-native conformations displayed correlations of around 0.6 or greater (33). Although there are still some inaccuracies in the predictions, these data suggest that SHAPER can provide reasonable predictions for structural features in native 23S RNA.

We then used SHAPER to compare the predicted and experimental reactivity of WT mini-PTC. The RNA tertiary structure used for SHAPER predictions was derived from an exceptionally high-resolution *E. coli* ribosome cryo-EM structure (PDB 7K00) (1). PDB 7K00 was modified prior to SHAPER analysis by deleting the small subunit, all large subunit r-proteins, and all 23S RNA residues save for residues 2043-2091, 2228-2258 and 2426-2625. The only addition to the structure was an AAAA tetraloop between residues A2092-A2095. Overall, the correlation between predicted and experimental SHAPE reactivity for WT mini-PTC is 0.61, suggesting that WT mini-PTC has near-native conformation that reasonably recapitulates most elements in the native ribosome.

To allow better comparison between two data sets, we applied normalization procedures to both predicted and experimental SHAPE data for WT mini-PTC, scaling the values within the range of 0 to 1 (**Figure 3A**). Differential SHAPE reactivity was calculated by subtracting the normalized predicted SHAPE reactivity from the normalized experimental counterpart. The differential SHAPE reactivity data was best fitted to the Cauchy probability density function using Fitter (**Figure S4**). We then applied the interquartile rule to identify extreme outliers as described in **Materials and Methods (Figure 3B**), which resulted in nine residues that exhibit differential SHAPE reactivities that either exceed 0.47 (colored red in **Figure 3C**) or fall below - 0.43 (colored blue in **Figure 3C**). These nine residues are located in four regions of WT mini-PTC including the L75-L80 homologation, L91, central loop, and L73-L93.

**Figure 3.**
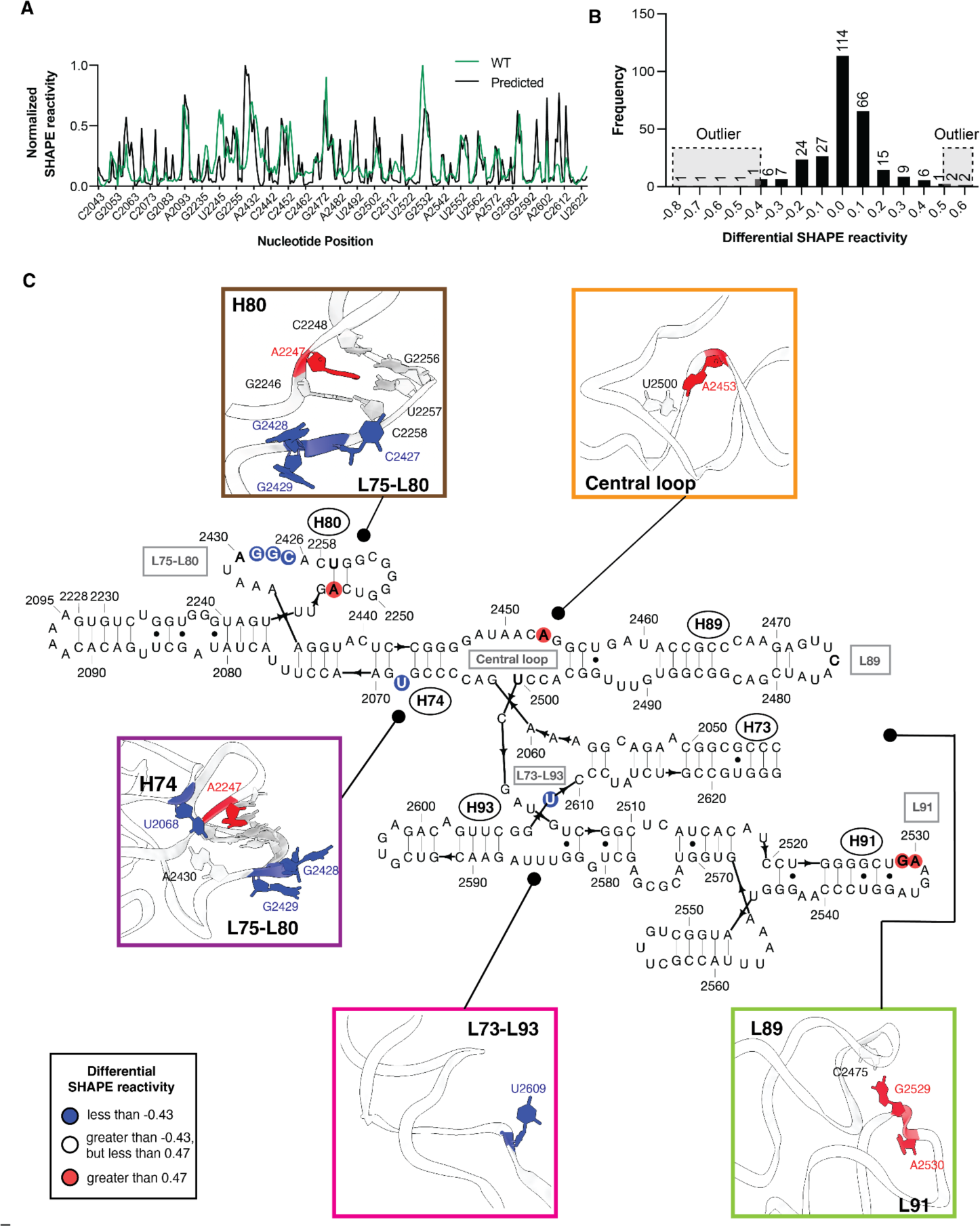
SHAPE analysis suggests that the folded structure of WT mini-PTC deviates from the canonical PTC fold in four important regions. **(A)** Normalized predicted SHAPE reactivity profile (black) and experimental SHAPE reactivity profile (green) of WT mini-PTC. **(B)** Histogram showing distribution of differential SHAPE reactivity between experimental data and predicted data of WT mini-PTC. Extreme outliers are highlighted with gray boxes and dashed black lines above histogram. Bin width of the histogram is equal to 0.1. Number of values in each bin were indicated above histogram. **(C)** Differential SHAPE reactivity between experimental data and predicted data was visualized on the predicted secondary and tertiary structure of WT mini-PTC. Extremely high and low outliers were plotted on the secondary structure as red and blue circles, respectively. Residues described in Results were bolded in the secondary structure diagram. Tertiary structures of highlighted WT mini-PTC regions were shown inside different color boxes, including L75-L80 (brown and violet), L89-L91 (light green), central loop (orange), and L73-L93 (pink).

Out of these nine residues, five were grouped in proximity to the L75-L80 homologation (brown box in **Figure 3C**). In this region C2427, G2428, and G2429 are predicted to form a loop and have high reactivity, while the experimental reactivity is low. In a similar way, A2247, located within H80, is predicted to pair with U2257 and thus show low reactivity, while the experimental reactivity is high. The fifth residue within this region that exhibits aberrant reactivity is located near H74 (purple box in **Figure 3C**). Specifically, U2068 is predicted to base-pair with A2430 but displays lower reactivity than anticipated (purple box in **Figure 3C**).

The remaining four residues within WT mini-PTC that display aberrant reactivity are located within the loop between H89 and H91, the central loop, and the loop between H73 and H93 (L73-L93). In the predicted structure, L89 and L91 engage in a loop-loop interaction facilitated by the base pairing of C2475 in L89 and G2529 in L91. As a result, these residues are predicted to display low SHAPE reactivity, while the experimental reactivity is high (light green box in **Figure 3C**). The remaining two residues exhibiting the highest deviations are situated within the central loop and L73-L93. A2453, positioned in the central loop, is predicted to possess low SHAPE reactivity due to its base-pair interaction with U2500; however, the experimental reactivity is high (orange box in **Figure 3B**). In L73-L93, a flexible residue U2609 demonstrates lower experimental reactivity than the predicted value (pink box in **Figure 3B**). Taken together, these results suggest that the WT mini-PTC sequence does not support a number of tertiary interactions that help define the overall shape of the PTC in the native *E. coli* ribosome. The greatest discrepancy is seen in the L75-L80 region, followed by L89-L91, the central loop, and L73-L93. Given these deficiencies, we turned to Eterna players to help identify changes in the WT mini-PTC sequence that would remedy these deficiencies and favor a structure that more closely resembles that of the native PTC.

### Crowdsourced design of diverse mini-PTC variants

We launched a unique Eterna challenge whose goal was to improve deficiencies in the WT mini-PTC fold, especially the interactions in the L75-L80 region, the loop between L89 and L91, L73 and L93, and within the central loop. In a typical Eterna challenge, players are tasked with designing *de novo* RNA sequences that fold into a target secondary structure. In this case, we asked players to identify mutations within a pre-established RNA sequence (WT mini-PTC) that would improve secondary and tertiary folding within a few key regions. As in all Eterna challenges, after designing sequences that fulfill the requirements of the challenge (DESIGN PHASE), players submit results and together rank the designs (VOTE PHASE). The top-ranked RNA designs are then synthesized and characterized by chemical probing methods to experimentally evaluate the extent to which each new design recapitulates the expected fold (ANALYZE PHASE) (**Figure 4A**).

**Figure 4.**
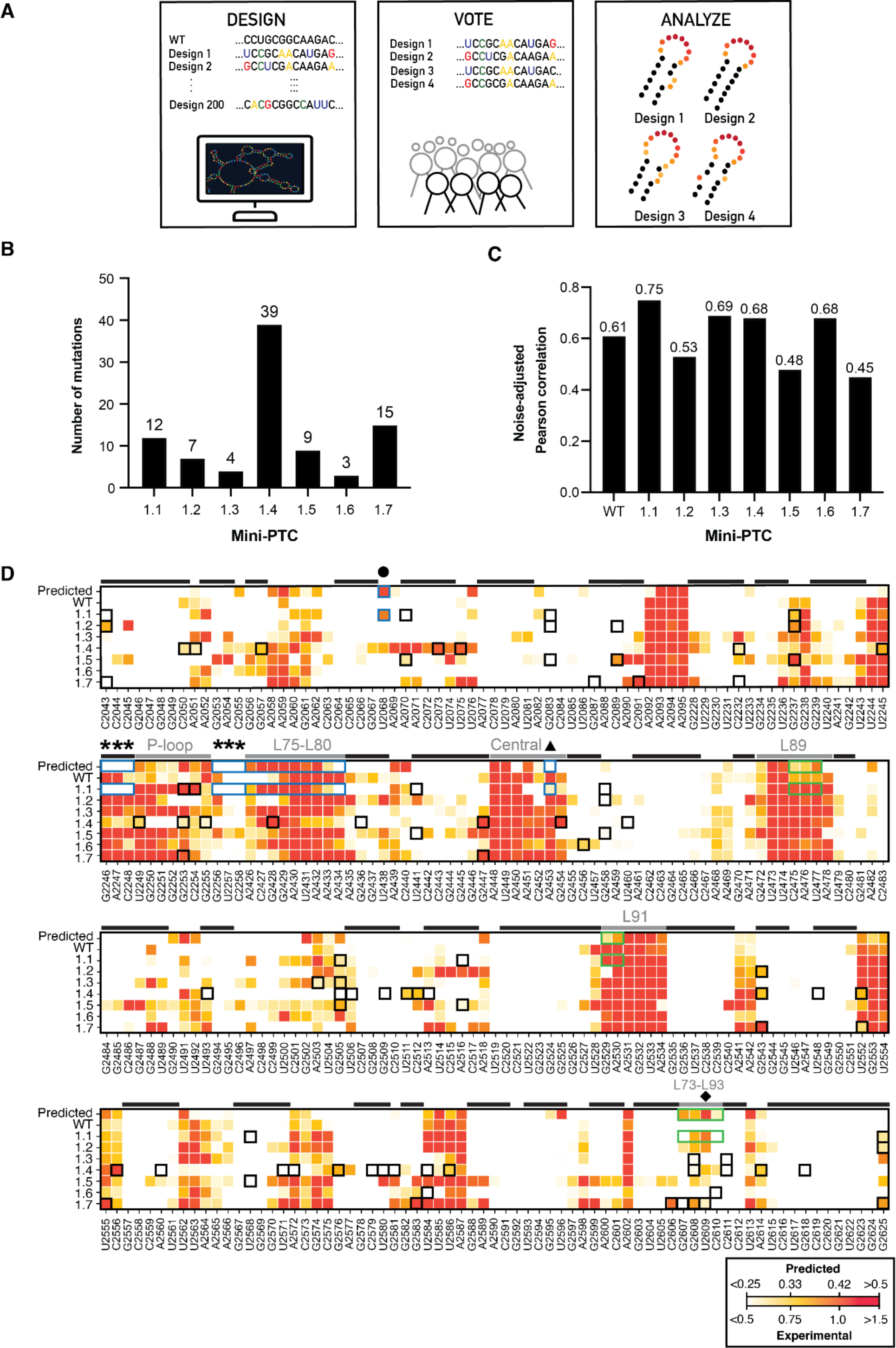
Eterna challenge allows selection of mini-PTC designs 1.1-1.7 that show diverse secondary structures. (**A)** Process in Eterna challenges for designing RNA molecules. Eterna players design new RNA constructs by making mutations within the provided sequence. Hundreds of designs were submitted by players at this stage. Arbitrary sequences are shown here for simplicity. Selected number of designs were evaluated by players voting and synthesized. Chemical probing methods were used to analyze the secondary structure between each design. **(B)** A bar graph summarizes the number of mutations of seven Eterna Mini-PTC. **(C)** A bar graph shows the noise-adjusted Pearson correlation between predicted SHAPE reactivity profile and experimental reactivity profile of WT mini-PTC and seven Eterna mini-PTC designs. (**D)** Heatmap showing the predicted and experimental SHAPE reactivity of the WT mini-PTC and seven Eterna mini-PTC designs from the first challenge (mini-PTC 1.1-1.7). SHAPE reactivity was plotted on a red-yellow-white spectrum where red represents high SHAPE reactivity (>0.5 predicted; >1.5 experimental) and white represents low SHAPE reactivity (<0.25 predicted; <0.5 experimental). Variations between experimental and predicted SHAPE reactivity were attributed to higher background in the experimental data. Base-paired residues and loops described in Results were marked with a black or gray line, respectively, over the heatmap. Residues were identified on the bottom using WT *E. coli* 23S rRNA numbering. Mutations in mini-PTC 1.1-1.7 were highlighted using a black square box. Residues in mini-PTC 1.1 that exhibited folding patterns closer to the predicted pattern were indicated using a blue box, whereas residues of mini-PTC 1.1 that exhibited differences from the prediction were indicated using a green box. Residues in H80 (G2246-C2248 and G2256-C2258), A2453, U2068, and U2609 were indicated using asterisks, triangle, circle, and diamond, respectively.

Several criteria provided boundary conditions for the WT mini-PTC challenge. Initially, we considered instructing players to limit their substitutions to only non-universally conserved nucleotides within WT mini-PTC. According to this plan, the only nucleotides in WT mini-PTC that could be varied during the challenge were those that varied in more than 1% of all gammaproteobacterial 23S sequences within the Comparative RNA Web (CRW) (7). This constraint would have severely limited the available mutational scope during the challenge, as only 44 of the 284 nucleotides in WT mini-PTC could be altered by any given player. To loosen the constraints, we allowed mutation at any position that previous data had shown retained at least 40% of wild type ribosome activity (45). This change increased the number of mutable bases from 44 to 79. The final instructions presented to players allowed them to substitute up to 40 of these 79 residues; their goal was to introduce mutations that would correct the folding deficiencies in WT mini-PTC. At the conclusion of this challenge, we received a total of 247 designs from 31 players. The top seven mini-PTC designs were ranked by player vote and termed mini-PTC 1.1-1.7. Examination of these sequences revealed significant diversity in the number and location of base changes introduced by any given player. Taken together, the top-ranked mini-PTC variants were diverse, with between 3 and 39 new mutations scattered throughout the mini-PTC sequence (**Figure 4B**, **Table 1, and Figures S5A-S11A**).

**Table 1.** Mutations introduced into WT mini-PTC in the seven top-ranked mini-PTC designs resulting from EteRNA challenge #1.

| **Name** | **# of mutations** | **Identity of Mutations** |
| --- | --- | --- |
| Mini-PTC 1.1 | 12 | C2043U, A2070C, G2083A, G2237A, G2253A, C2254U, U2441G, G2458A, G2505U, A2516G, U2568C, G2625A |
| Mini-PTC 1.2 | 7 | C2043U, G2083A, C2089U, G2237A, G2458A, G2543A, G2625A |
| Mini-PTC 1.3 | 4 | A2503U, G2505U, G2608A, C2611U |
| Mini-PTC 1.4 | 39 | C2050U, A2051U, G2057C, C2073G, U2075A, C2232U, U2245A, U2249A, G2253A, G2255A, G2428A, G2436C, G2447A, G2454A, U2460A, U2493A, G2505A, U2506C, G2509U, U2511A, C2512A, A2513G, G2543A, U2548A, U2552A, C2556A, A2560U, U2571C, A2572G, G2576A, C2579A, U2580C, G2581U, U2584C, U2586A, G2608A, C2611G, A2614U, G2618A |
| Mini-PTC 1.5 | 9 | A2070C, G2083A, C2089U, G2237A, U2441G, G2458A, G2505U, A2516G, U2568C |
| Mini-PTC 1.6 | 3 | C2456U, U2584A, C2610G |
| Mini-PTC 1.7 | 15 | C2043U, G2087A, C2091U, C2232U, G2253A, G2447A, G2543A, U2552A, U2555A, G2583A, C2606U, G2607C, G2608A, U2609A, G2625A |

### Mini-PTC 1.1 improves folding into a native PTC structure

With the seven new mini-PTC sequences in hand, we turned to SHAPE analysis to determine if the introduced mutations alleviated the folding deficiencies identified in the Eterna challenge. All seven Eterna player-designed mini-PTCs (mini-PTC 1.1-1.7) were prepared by run-off transcription and purified to homogeneity (**Figure S1A**). Native gel analysis revealed that all the products were homogeneous when refolded at at 50 °C in the presence of 10 mM MgCl_2_ (**Figure S12**). These conditions were then used for SHAPE analysis (**Figure S5B-S11B**) and the correlation between noise-adjusted experimental SHAPE reactivity and SHAPER-predicted reactivity was established. The seven Eterna-designed mini-PTC sequences were classified into two categories depending on whether this correlation was better or worse than that of WT mini-PTC (correlation = 0.61). We found that mini-PTC 1.2, 1.5, and 1.7 exhibited lower correlations with scores of 0.53, 0.48, and 0.45, respectively, while mini-PTC 1.3, 1.4, and 1.6 were improved with scores of 0.69, 0.68, and 0.68, respectively **(Figure 4C)**. Out of the seven Eterna mini-PTC designs, mini-PTC 1.1 exhibited the highest correlation with a score of 0.75; this sequence was thus chosen for more detailed study.

To better understand how mini-PTC 1.1 outcompetes other designs in its ability to recapitulate the native PTC fold, we generated a heatmap showing the predicted and experimental SHAPE reactivities of WT mini-PTC and all seven Eterna mini-PTC designs (**Figure 4D**). We then analyzed the SHAPE reactivities of the Eterna mini-PTC designs in the four regions previously identified as having folding deficiencies in the WT mini-PTC. Notably, mini-PTC 1.1 exhibits significant improvement in two of these regions - the central loop and the L75-L80 junction near the P-loop (blue boxes in **Figure 4D**). In the central loop, only mini-PTC 1.1 shows low SHAPE reactivity at residue A2453 that aligns with the low predicted SHAPE reactivity, whereas all other mini-PTC designs display high reactivity at this position (triangle in **Figure 4D**). In H80, only mini-PTC 1.1 shows the expected low SHAPE reactivities at the 2246:2258, 2247:2257, 2248:2256 base pairs that match with prediction (asterisk in **Figure 4D**). Adjacent to H80, SHAPE reactivity of mini-PTC 1.1 in L75-L80 agrees well with the predicted SHAPE reactivity (L75-L80 in **Figure 4D**). Residue U2068, which previously exhibits low reactivity in WT mini-PTC, has relatively high SHAPE reactivity in mini-PTC 1.1 that is similar to the predicted SHAPE reactivity (circle in **Figure 4D**). However, differences between the experimental and predicted SHAPE reactivity of mini-PTC 1.1 were observed in the L89-L91 and L73-L93 (green boxes in **Figure 4D**). Residues C2475-U2477 in L89 and residues G2529-A2530 in L91 display higher reactivities than the prediction (L89 and L91 in **Figure 4D**). In addition, mini-PTC 1.1 exhibits a relatively low SHAPE reactivity pattern in L73-L93, including U2609, compared to the predicted SHAPE reactivities (diamond in **Figure 4D**). While this analysis indicates that mutations in mini-PTC 1.1 demonstrate substantial enhancement in the central loop, H80, and L75-L80 over WT mini-PTC, they also suggest some remaining deficiencies with respect to interactions in the L89-L91 and L73-L93.

### Mini-PTC 1.1 binds ribosome substrate analogs

During translation, the ribosomal P site is occupied by a tRNA whose 3′-ribose hydroxyl group is acylated by the growing peptide chain, whereas the A site is occupied by a tRNA whose 3′-ribose hydroxyl group is acylated by the incoming α-amino acid. Several substrate mimics have been devised to enable single-turnover analysis of peptide bond formation by purified ribosomes and variants thereof (46). A widely used mimic of the P-site peptidyl-tRNA is CC-puromycin-phenylalanine-caproic acid-biotin (CCA-pcb), and a widely used mimic of the A-site acyl-tRNA is C-puromycin (C-pmn), in which a non-hydrolyzable amide bond ensures single-turnover reactivity (**Figure 5A**) (47, 48). We made use of CCA-pcb and C-pmn in combination with SHAPE to determine whether these substrate mimics interact with WT mini-PTC and mini-PTC 1.1.

**Figure 5.**
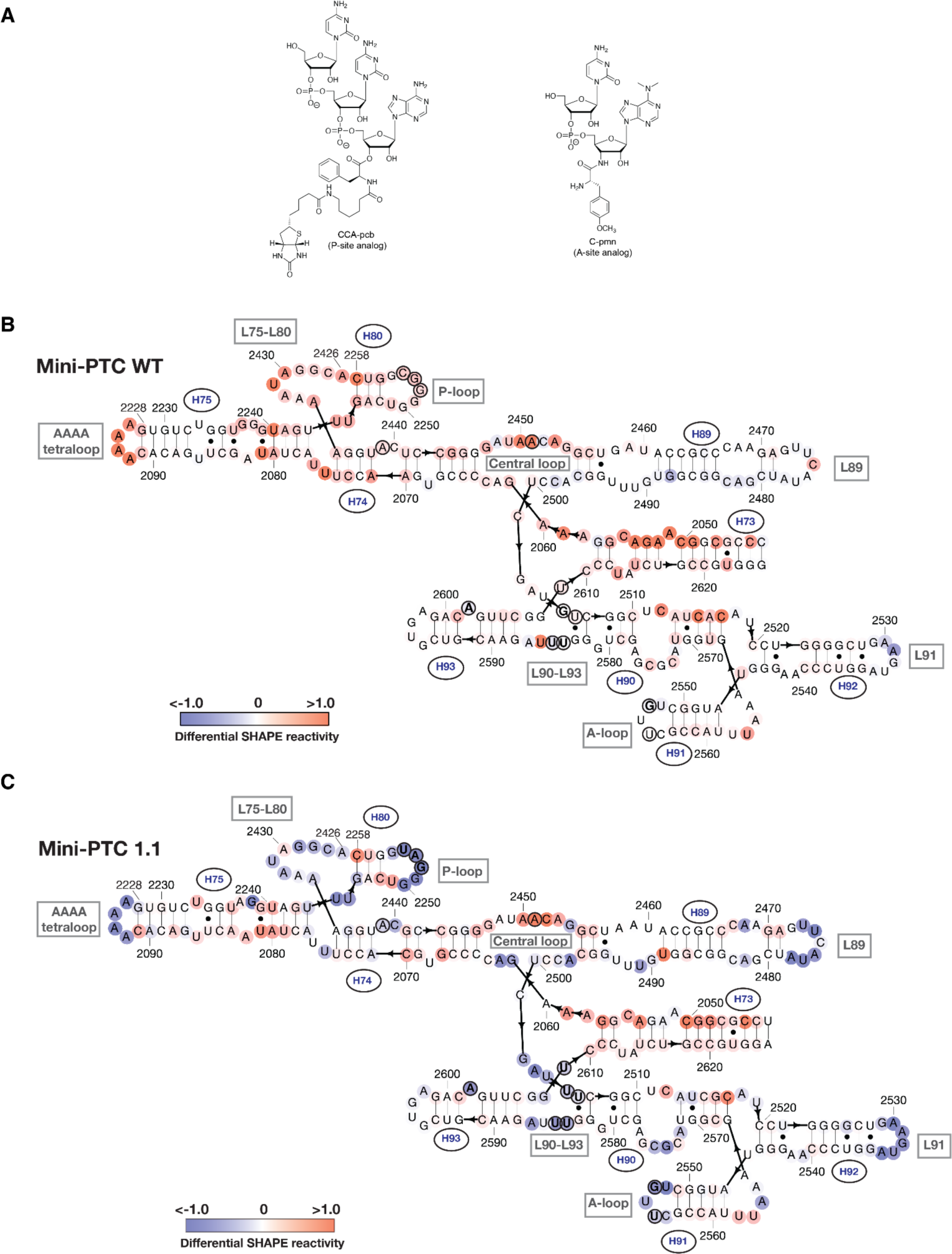
Mini-PTC 1.1 engages the ribosomal A- and P-site substrate mimics better than WT mini-PTC **(A)** Chemical structures of CCA-pcb (P-site analog) (left), and C-pmn (A-site analog) (Right). **(B-C)** Predicted secondary structure diagram showing differential SHAPE reactivities of WT mini-PTC **(B)** and mini-PTC 1.1 **(C)** in the presence of 100 µM CCA-pcb and 100 µM C-pmn. Differential SHAPE reactivities were represented on a blue-white-red spectrum, with blue indicating low changes, red indicating high changes, and white indicating no changes in SHAPE reactivity upon substrate binding. Previously identified 13 PTC residues that become protected upon binding A- or P-site tRNA substrates in the ribosome-tRNA complexes were indicated in circled residues. 5 of 13 residues in WT mini-PTC **(B)** and 12 of 13 residues in mini-PTC 1.1 **(C)** that become protected upon binding substrate mimics were highlighted as bold residues.

We performed SHAPE analysis of WT mini-PTC and mini-PTC 1.1 in the absence or presence of 100 µM or 200 µM of both CCA-pcb (P-site substrate analog) and C-pmn (A-site substrate analog) (**Figure S13**). Examination of the SHAPE reactivity as a function of nucleotide reveals that many positions within WT mini-PTC become more reactive towards 1M7 as the concentration of CCA-pcb and C-pmn increases from 0 to 200 µM, with the most substantial changes at positions corresponding to nucleotides A2093 (within the AAAA tetraloop), G2472 (within the loop that caps H89), G2532 (within the loop that caps H91), and between U2245 and U2255 (within the P-loop). In contrast, many positions within mini-PTC 1.1 become less reactive towards 1M7 as the concentration of CCA-pcb and C-pmn increases from 0 to 200 µM. Although the magnitude of the the changes are more modest, statistically significant reductions in reactivity are observed between U2245 and U2255 (in the P-loop), between U2472 and U2482 (within the loop that caps H89), and near G2532 (within the loop that caps H91). The locations of changes in reactivity detected at 100 µM CCA-pcb and C-pmn are mapped on the native PTC secondary structure in **Figure 5B and C**. Nucleotides that become less reactive (more protected) upon the binding of substrates are represented in blue, while those that become more reactive (less protected) are shown as red.

More fine-grained analysis of the SHAPE reactivity changes upon addition of CCA-pcb and C-pmn confirms that the nucleotide substitutions embedded into mini-PTC 1.1 during the Eterna challenge not only improve its ability to fold into a PTC-like structure, but also its ability to bind PTC substrates. In particular, the SHAPE reactivity of mini-PTC 1.1 in the presence of CCA-pcb and C-pmn is reduced substantially within the P-loop and the A-loop, as well as within L89, L91, and the AAAA tetraloop (**Figure 5C**). These decreases in SHAPE reactivity are not observed when CCA-pcb and C-pmn were added to WT mini-PTC (**Figure 5B**). To investigate whether mini-PTC 1.1 binds CCA-pcb and C-pmn in the appropriate regions, we compared its differential SHAPE reactivity to that obtained previously for intact wild-type ribosome-tRNA complexes (49). These previous analyses identified 13 PTC residues that become protected upon binding A- or P-site tRNA substrates (circled residues in **Figure 5B and C**) (49). Twelve of these thirteen positions within mini-PTC 1.1 show more dramatically reduced SHAPE reactivity in the presence of 100 µM CCA-pcb and C-pmn (bold residues in **Figure 5C**), whereas only WT mini-PTC reveals that 5 of the 13 residues of WT mini-PTC show reduced SHAPE reactivity, and the reduction is modest (bold residues in **Figure 5B**).

### Effects of Mini-PTC 1.1 on the rate of peptide bond formation

The previous results presented herein provide evidence that mini-PTC 1.1 associates with substrate analogs CCA-pcb and C-pmn when present at 100 µM concentration and reorganizes substantially upon binding in a manner consistent with the expected tertiary fold of the PTC. We next asked whether this association could promote a bond-forming reaction between CCA-pcb and C-pmn to generate C-puromycin-phenylalanine-caproic acid-biotin (C-pmn-pcb) in a variation of the well-established “fragment reaction” (**Figure 6A**) (46, 47). CCA-pcb and C-pmn (100 µM each) were added to mini-PTC 1.1 (20 µM) or 50S ribosomes (0.1 µM) and the formation of C-pmn-pcb was analyzed as a function of incubation time and temperature. Rather than detecting C-pmn-pcb directly, we treated it with RNase A and monitored formation of the so-generated puromycin-phenylalanine-caproic acid-biotin (pmn-pcb) digestion product using LC-MS. Incubations were performed at 4 °C and 50 °C as well as at −20 °C, as previous reports have shown that reaction at eutectic conditions could enhance polymerase ribozyme activity, as the cold temperature better reflects the conditions in the primordial RNA world (50). To perform quantitative analysis, activity was calculated by a ratio of the peak area corresponding to pmn-pcb (**Figure S14**) to the peak area of co-injected Leu-enkephalin standard, then multiplying the ratio by 10^3^ (**Figure S15** and **Table S1, S2**).

**Figure 6.**
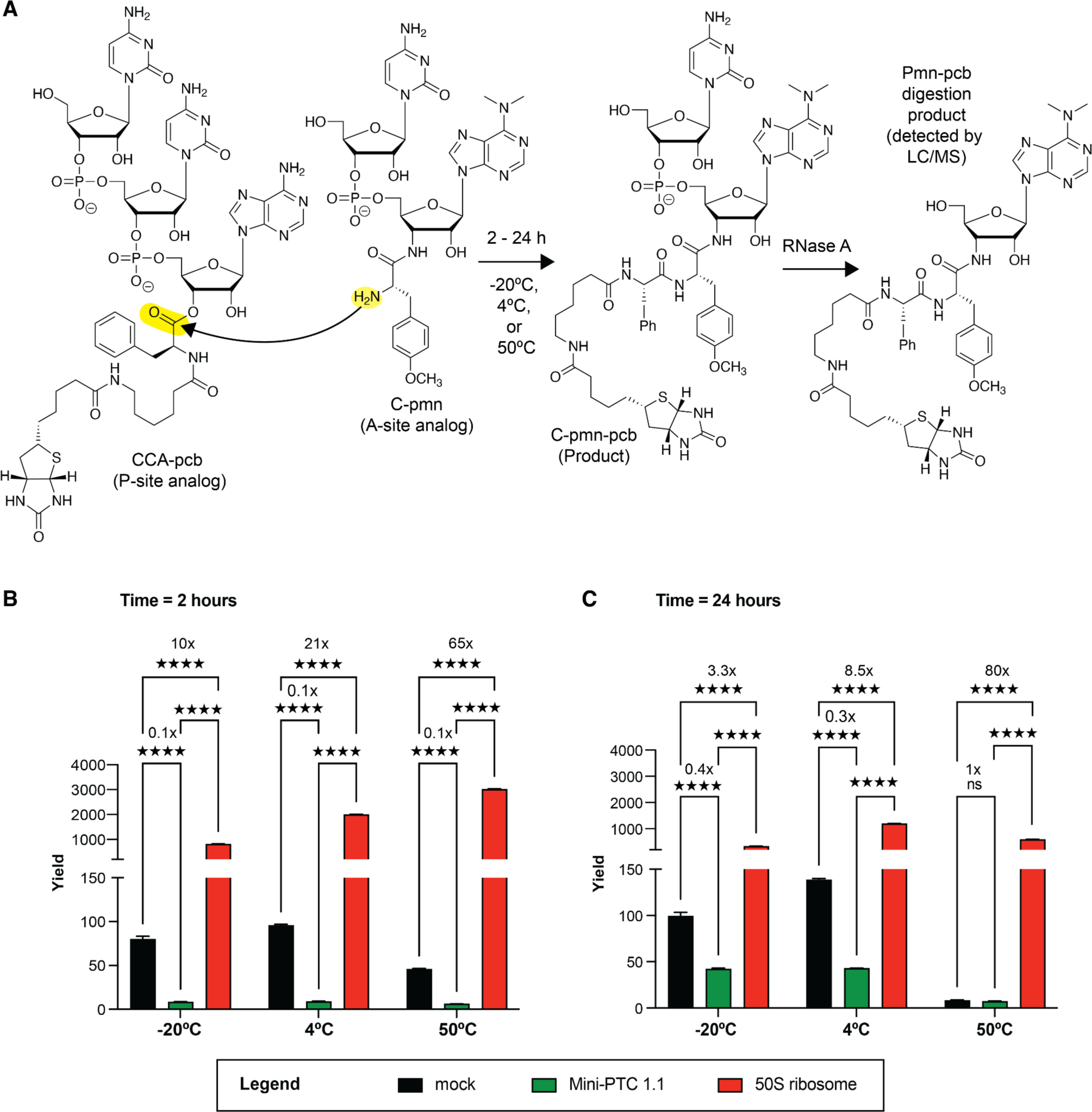
Mini-PTC 1.1 inhibits the peptidyl transferase activity as evaluated by the fragment reaction. **(A)** Schematic representation of fragment reaction, followed by RNase A/LC-MS assay. CCA-pcb and C-pmn were used as a P-site and an A-site analog, respectively. The reaction was incubated with varied time (2 or 24 h) and temperature (−20 °C, 4 °C or 50 °C). The product from this reaction (C-pmn-pcb) was subsequently digested by RNase A that cleaves off terminal puromycin-pcb (pmn-pcb), which can be quantified by LC-MS analysis. Leu-enkephalin was used as an internal standard. Reactions were run with water (mock), 20 µM mini-PTC 1.1, and 0.1 µM *E. coli* 50S ribosomal subunit at −20°C, 4°C, or 50°C for 2 hours **(B)** or 24 hours **(C).** Reactions were run as triplicates (n = 3). The yield of the fragment reaction between water (mock), mini-PTC 1.1, and *E. coli* 50S ribosomal subunit was compared by using two-way ANOVA with Tukey’s multiple comparison test. \*\*\*\**p*≤ 0.0001, \*\*\**p* ≤ 0.002,\*\**p* ≤ 0.021, \**p* ≤ 0.032, ns ≤ 0.1234.

Co-incubation of CCA-pcb and C-pmn with 50S ribosomes led to rapid formation of C-pmn-pcb products within 2 h, with little or no change after 24 h incubation (**Figure 6B, C**). Interestingly, although the overall yield of C-pmn-pcb generated using WT *E. coli* 50S subunits was slightly temperature-dependent, the yield relative to the yield in a mock reaction that lacked 50S subunits was highly temperature-dependent, with the highest increases observed at 50 °C and after both 2 and 24 h incubation. In contrast, co-incubation of CCA-pcb and C-pmn with mini-PTC 1.1 results in significantly less product than the mock reaction regardless of temperature and incubation time. Little or no C-pmn-pcb product was observed after 2 h incubation; after 24 h some product is formed at −20 °C and at 4 °C, but in most cases the yield was 3- to 10-fold lower than that observed in the absence of mini-PTC 1.1. Rather than promoting the reaction of CCA-pcb and C-pmn to generate an amide bond in C-pmn-pcb, mini-PTC 1.1 acts to inhibit the reaction. Taken with chemical probing analysis that supports the association of mini-PTC 1.1 with CCA-pcb and C-pmn, these results imply that the substrates are misoriented, which interferes with the peptide bond formation.

## DISCUSSION

The ribosome is a complex biological machine whose evolutionary conserved active site promotes rapid and specific amide bond formation. The mass of the *E. coli* ribosome, including both RNA and all r-proteins, tops out at 2.8 MDa. But the PTC–where bond formation occurs– constitutes only ∼3-4% of the total ribosome mass. The remaining mass serves many equally conserved functions–in addition to peptide bond formation within the PTC, the ribosome must assemble faithfully on an mRNA transcript, bind and accommodate acyl- and peptidyl-tRNAs, translocate these tRNAs after bond formation, thread a growing polypeptide chain through the exit tunnel, and ultimately catalyze peptidyl-tRNA hydrolysis once peptide synthesis is complete. All these steps must be performed rapidly and with exceptional fidelity. Attempts to recapitulate the structure, substrate recognition, and bond formation of an intact ribosome in the context of minimal PTC provide an opportunity to more directly understand the local factors that contribute to bond formation and inform current efforts to design ribosomes with altered function.

Several studies have been conducted with the goal of creating a minimal RNA construct derived from 23S rRNA that exhibits peptidyl transferase activity. Strobel and coworkers generated a 314-nt RNA construct derived from *B. stearothermophilus* 23S rRNA domain V that contain essential features of the PTC, including central loop, A-loop, and P-loop, but their minimal 23S rRNA construct did not possess catalytic peptidyl transferase activity even after multiple rounds of selection (51). More recently, several small rRNA subdomains derived from the 23S rRNA have been studied and showed ability to promote peptide bond formation between A- and P-site substrates (16, 18, 19). However, despite these findings, structural studies of 23S rRNA minimal constructs are not yet available to help define the molecular basis for their function. In this study, we used the Eterna citizen platform to improve folding of WT mini-PTC, which resulted in the identification of mini-PTC 1.1. Although Eterna does not explicitly evaluate tertiary interactions, subsequent SHAPE analysis suggests the mutations introduced in mini-PTC 1.1 led to improvements in tertiary folding. This improvement also translated to improved binding of A- and P-site substrate analogs. Although mini-PTC 1.1 showed improved tertiary folding and substrate binding relative to WT mini-PTC, it still has limitations in positioning substrates correctly as evidenced by the inhibition observed in the fragment reaction under most tested conditions. More in-depth structural studies are necessary to evaluate how mini-PTC 1.1 needs to be revised to promote a conformation suitable for catalyzing the peptide bond formation (52).

In summary, here we report the design of mini-PTC constructs and characterization of mini-PTC 1.1 RNA that recapitulates the structural features of the *E. coli* ribosomal peptidyl transferase center (PTC). Our work highlights the previously demonstrated value of the Eterna platform for robust design of RNAs with desired secondary and tertiary structures and elucidates the relationship between improved tertiary folding of a minimal 23S rRNA construct and its ability to bind A- and P-site substrates.

## DATA AVAILABILITY

All MATLAB and python scripts used for SHAPE analysis and determining outliers are available at https://github.com/gem-net/miniPTC-paper. Raw capillary electrophoresis data and data obtained from SHAPER predictions are available at **Supplementary Data.**

## ACKNOWLEDGEMENTS

The authors are grateful to members of the Schepartz, Das, and Cate labs for helpful discussion, and especially to Dr. Chandrima Majumdar, Cameron Swenson, Dr. Carly Schissel, Dr. Josh Walker, and Dr. Riley Fricke for comments on the manuscript. We thank J. Romano for expert administration of Eterna. We also thank Kristina Boyko (Cate lab) for providing purified *E. coli* 50S ribosomes, and Kittithat Krongchon (University of Illinois, Urbana-Champaign) for revising python scripts used to generate a heatmap of SHAPE results.

