## Supplementary Data for "Minimization of the E. coli ribosome, aided and optimized by community science"

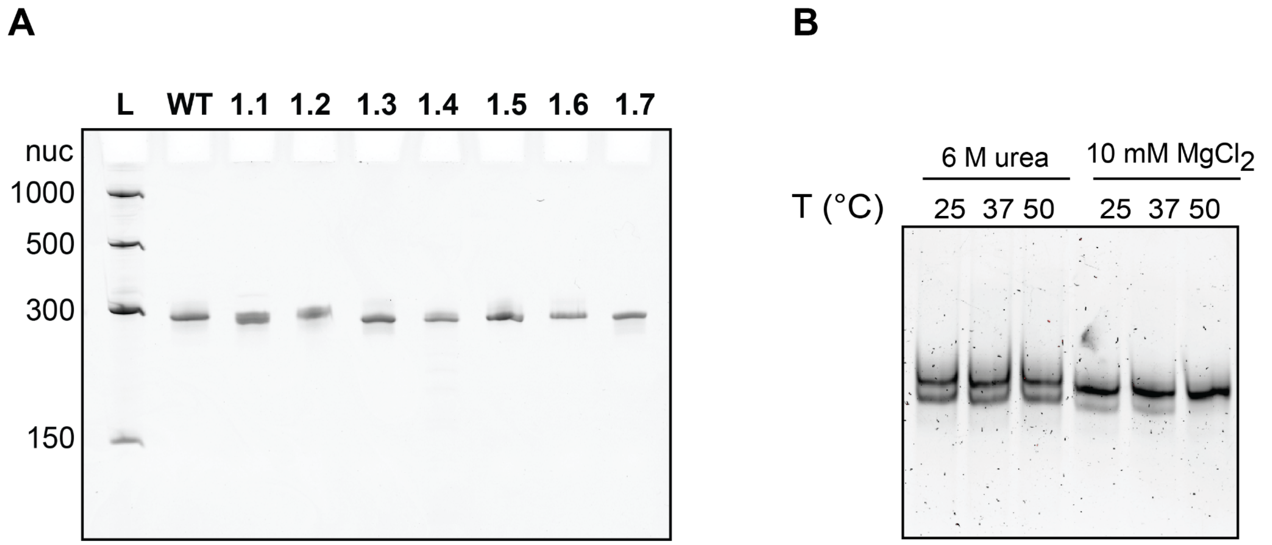


**Figure S1.** Native and denaturing gel electrophoretic analysis of WT mini-PTC and Eterna-generated designs mini-PTC **1.1-1.7**. **(A)** Purified samples of WT mini-PTC and mini-PTC **1.1-1.7** migrate as single bands on a 5% PAGE gel containing 7 M urea. The ladder used as a reference contains single stranded RNAs with 150, 300, 500, and 1000 nt. **(B)** Purified samples of WT mini-PTC were dissolved in 50 mM Na-HEPES, pH 8.0 supplemented with 6 M urea (lanes 1-3) or 10 mM $MgCl_2$ (lanes 4-6) and incubated at 25, 37, and 50 ºC for 20 min and applied to a 8% native acrylamide gel. WT mini-PTC migrated as a single band when re-folded in the presence of 10 mM $MgCl_2$ at 50 ºC.

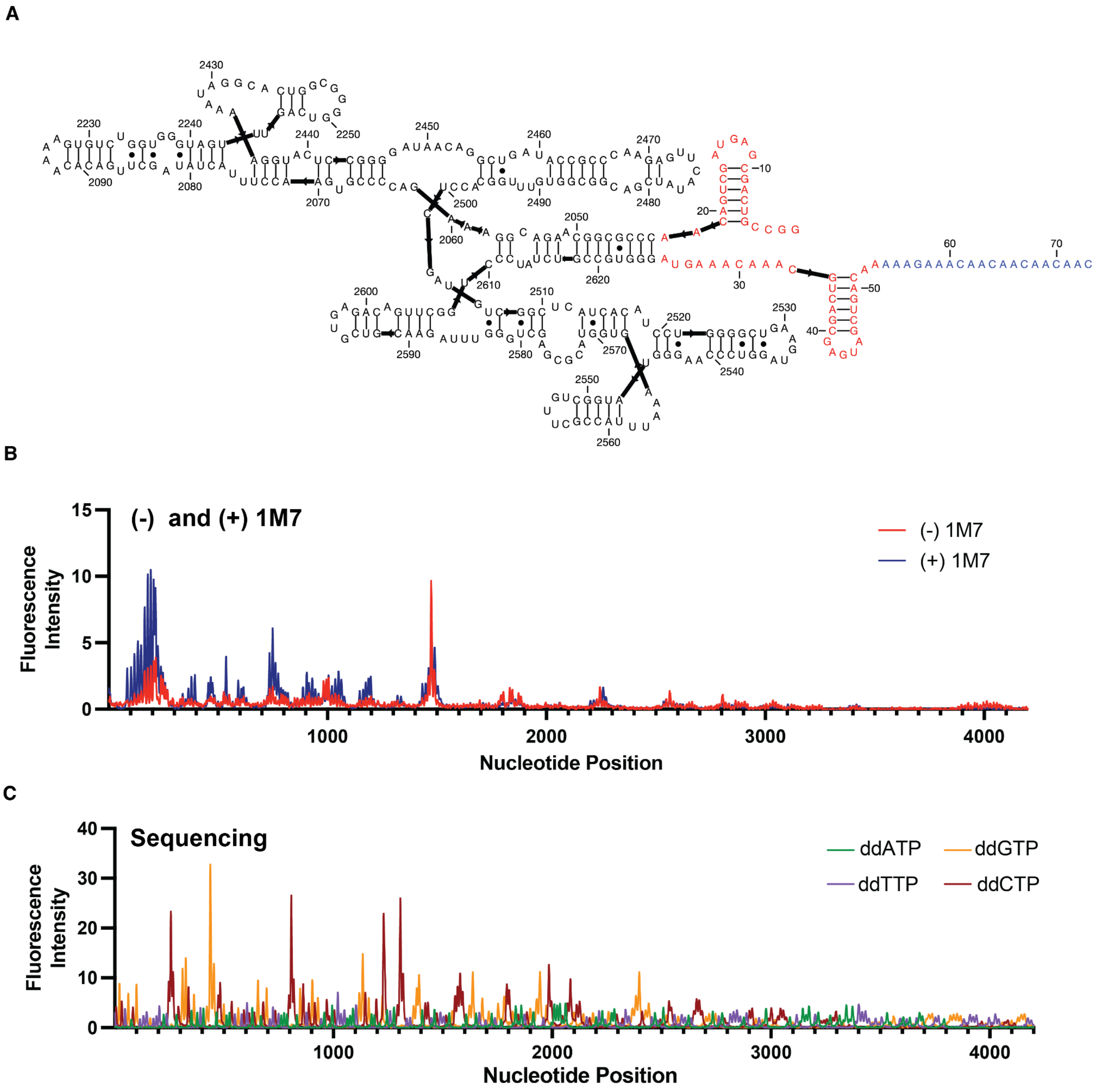


**Figure S2.** Design and capillary electrophoresis data of WT mini-PTC for SHAPE analysis. **(A)** The sequence of WT mini-PTC (284 nucleotides) drawn according to the secondary structure seen in the context of an intact ribosome. GAGUA reference hairpins (1) at 5′ and 3′ ends used to normalize SHAPE reactivity are shown in red. The sequence of the primer binding site is shown in blue. **(B)** Representative capillary electrophoresis trace illustrating the relative abundance of cDNA reaction products resulting from reverse-transcription of 1M7-treated (blue) or untreated (red) samples of WT mini-PTC. Peaks represent positions within WT mini-PTC that react significantly with 1M7; the peak in the untreated sample represents a position in the sequence that naturally impedes the processivity of reverse transcriptase. **(C)** Sequencing ladder used to assign nucleotide positions of WT mini-PTC; ddATP (green), ddTTP (purple), ddGTP (yellow), and ddCTP (red).

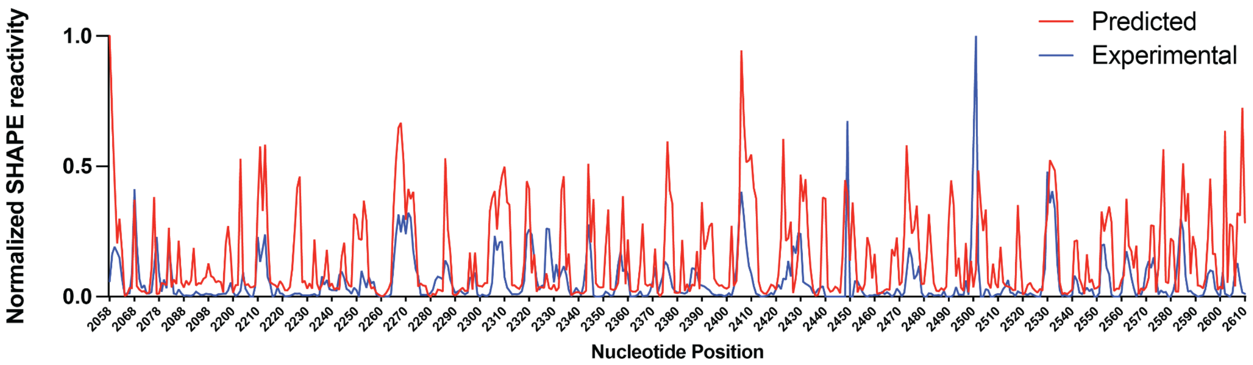


**Figure S3.** SHAPER can reasonably predict reactivity with good agreement in the context of 23S RNA. The normalized predicted SHAPE reactivity profile (red) and the experimental SHAPE reactivity profile (blue) of domain V nucleotides obtained from a previous report by Deigan and coworkers (2).

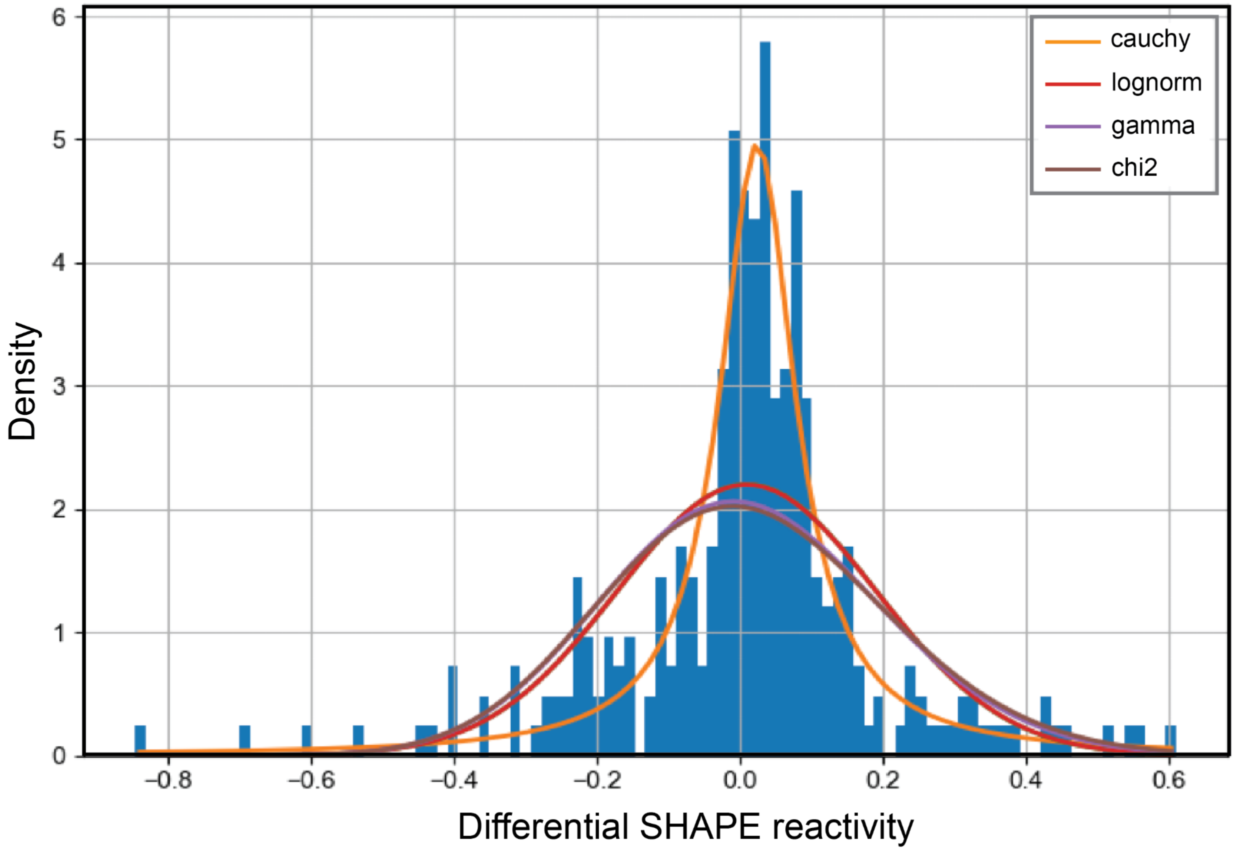


**Figure S4.** Differential SHAPE reactivity between predicted and experimental SHAPE was fitted into various types of distributions using Fitter. The differential SHAPE reactivity of WT mini-PTC data, represented by the blue histogram, was best fitted to the Cauchy probability density function (orange line), followed by the log-normal (red line), gamma (violet line), and chi-squared probability density functions (brown line).

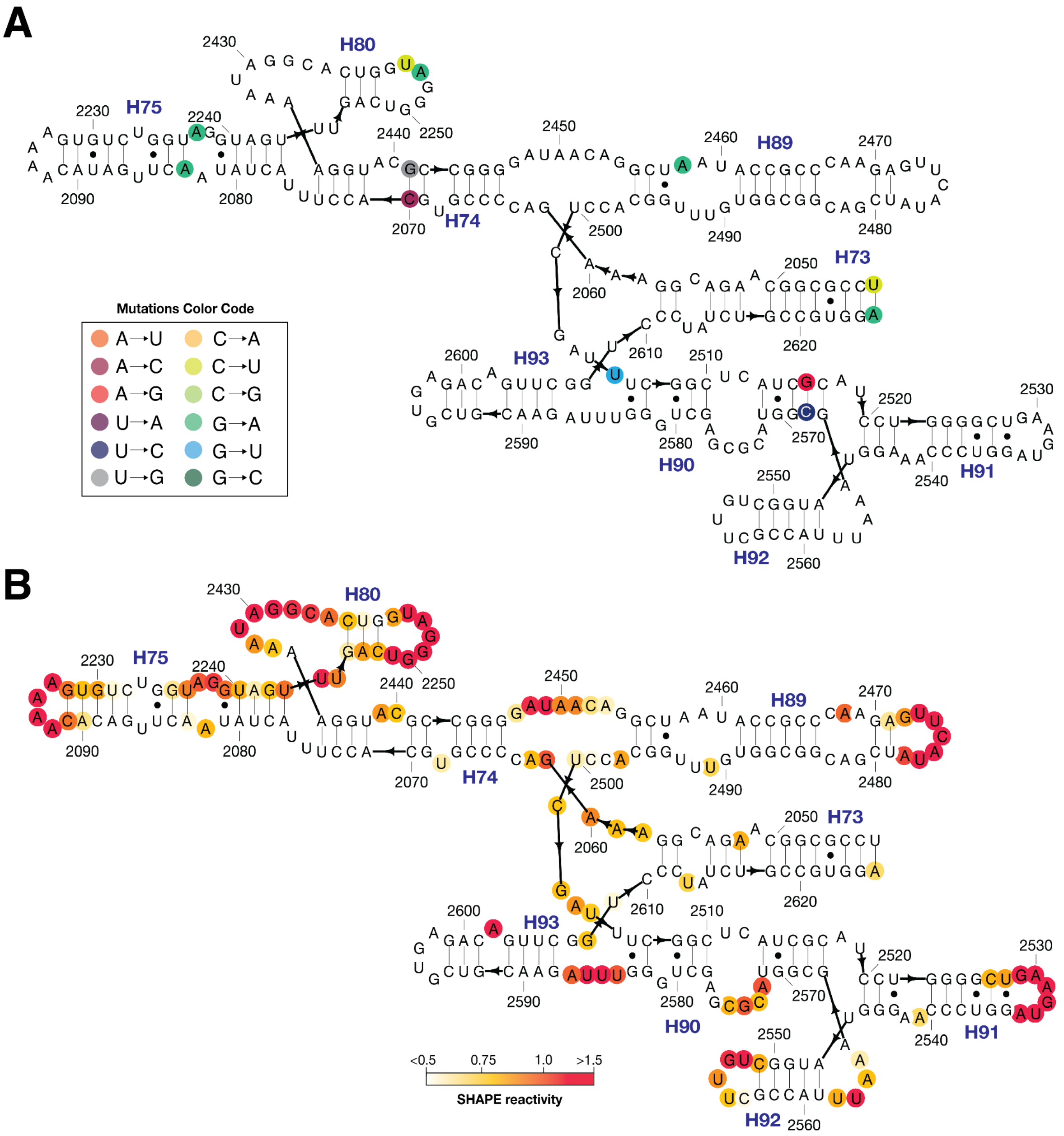


**Figure S5.** Predicted secondary structure and SHAPE analysis of mini-PTC 1.1. **(A)** Predicted secondary structure of mini-PTC 1.1. Residue numbers and helices are provided in black and blue. Mutations are highlighted in colored circles, which include mutations C2043U, A2070C, G2083A, G2237A, G2253A, C2254U, U2441G, G2458A, G2505U, A2516G, U2568C, and G2625A. **(B)** SHAPE analysis of mini-PTC 1.1. SHAPE reactivity was plotted in the red-yellow-white spectrum where red represents high SHAPE reactivity (>1.5) and white represents low SHAPE reactivity (<0.5).

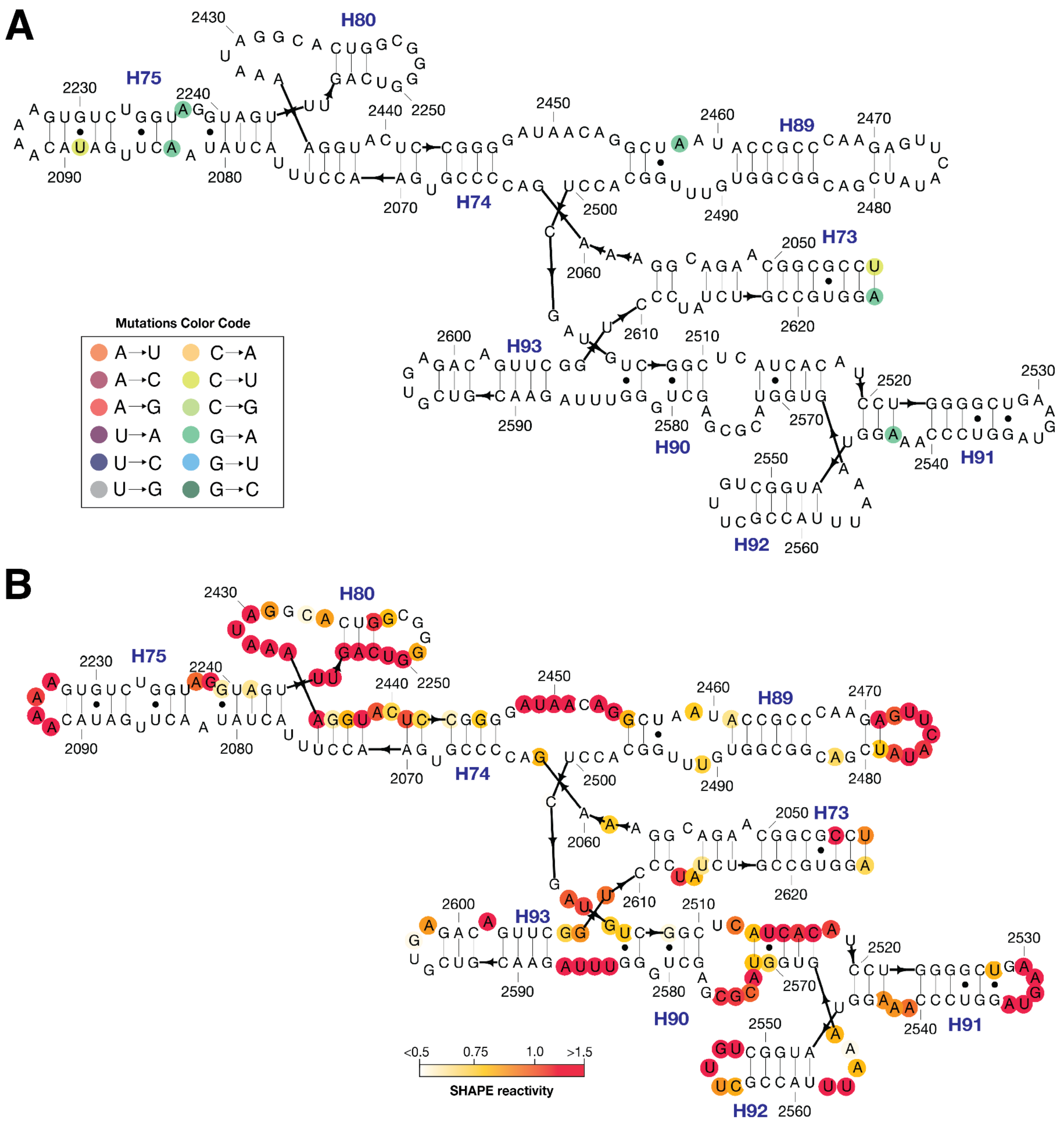


**Figure S6.** Predicted secondary structure and SHAPE analysis of mini-PTC 1.2. **(A)** Predicted secondary structure of mini-PTC 1.2. Residue numbers and helices are provided in black and blue. Mutations are highlighted in colored circles, which include mutations C2043U, G2083A, C2089U, G2237A, G2458A, G2543A, and G2625A. **(B)** SHAPE analysis of mini-PTC 1.2. SHAPE reactivity was plotted in the red-yellow-white spectrum where red represents high SHAPE reactivity (>1.5) and white represents low SHAPE reactivity (<0.5).

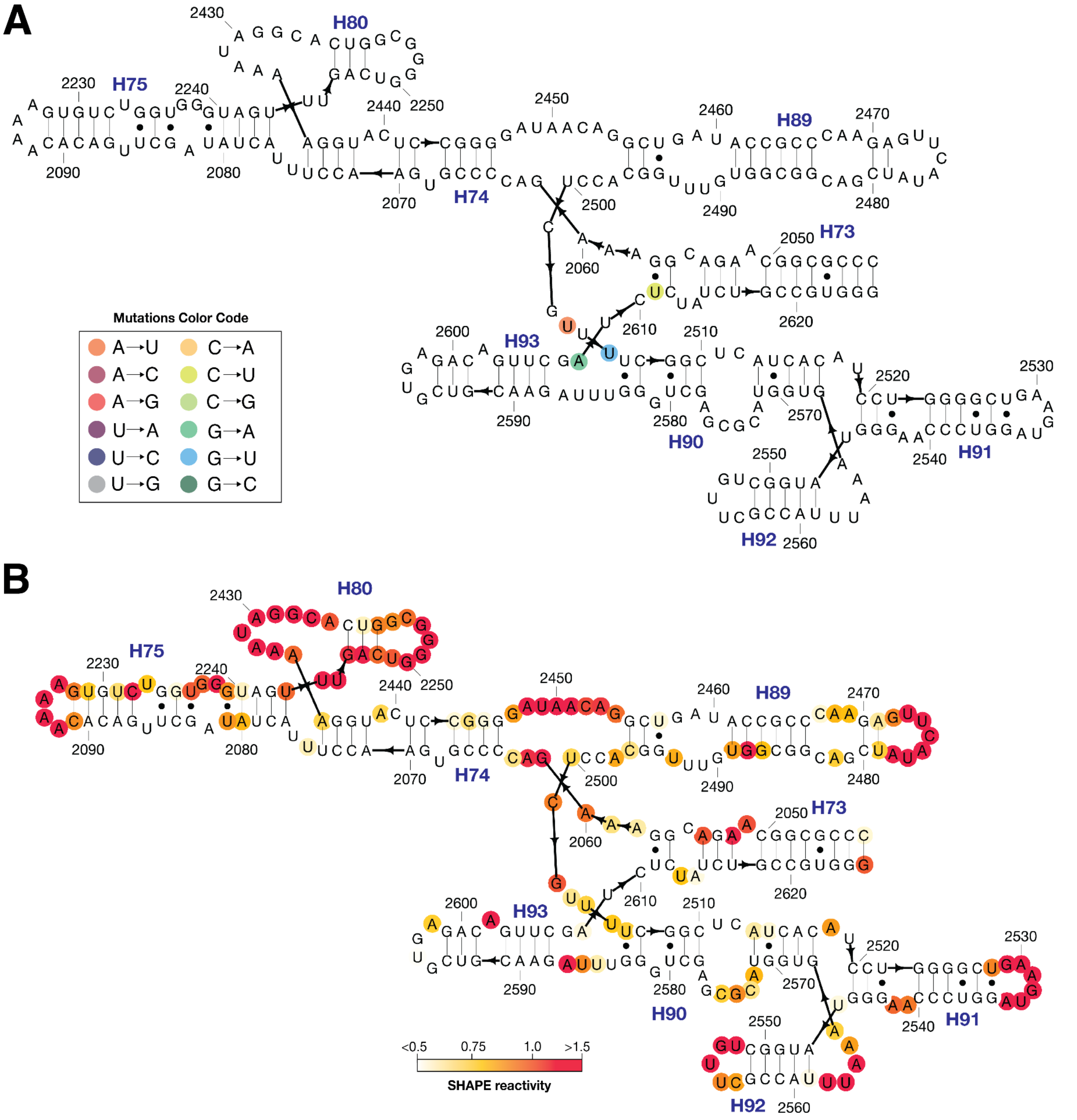


**Figure S7.** Predicted secondary structure and SHAPE analysis of mini-PTC 1.3. **(A)** Predicted secondary structure of mini-PTC 1.3. Residue numbers and helices are provided in black and blue. Mutations are highlighted in colored circles, which include mutations A2503U, G2505U, G2608A, and C2611U. **(B)** SHAPE analysis of mini-PTC 1.3. SHAPE reactivity was plotted in the red-yellow-white spectrum where red represents high SHAPE reactivity (>1.5) and white represents low SHAPE reactivity (<0.5).

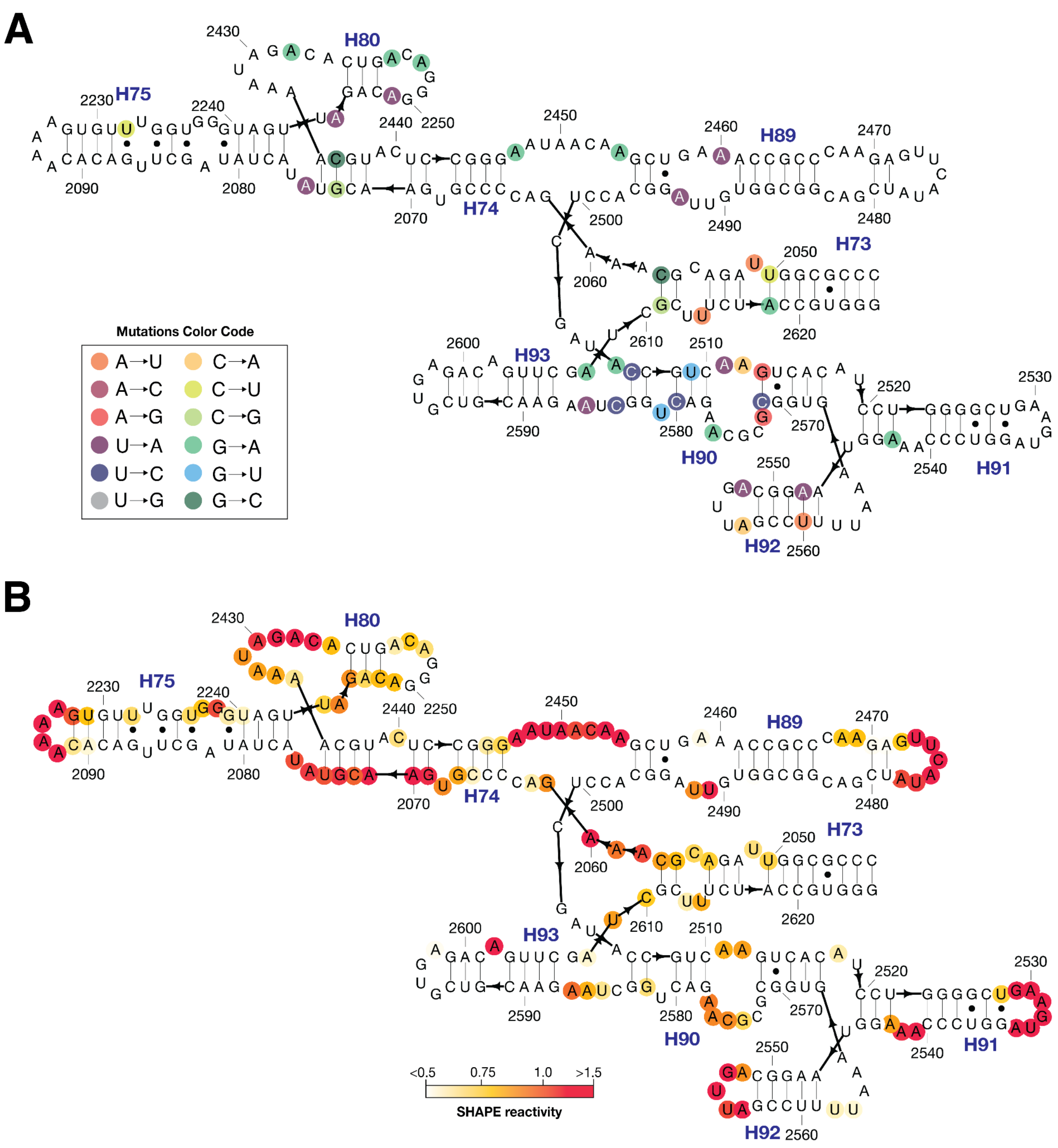


**Figure S8.** Predicted secondary structure and SHAPE analysis of mini-PTC 1.4. **(A)** Predicted secondary structure of mini-PTC 1.4. Residue numbers and helices are provided in black and blue. Mutations are highlighted in colored circles, which include mutations C2050U, A2051U, G2057C, C2073G, U2075A, C2232U, U2245A, U2249A, G2253A, G2255A, G2428A, G2436C, G2447A, G2454A, U2460A, U2493A, G2504U, U2506C, G2509U, U2511A, C2512A, A2513G, G2543A, U2548A, U2552A, C2556A, A2560U, U2571C, A2572G, G2576A, U2580C, G2581U, U2584C, U2586A, G2608A, C2611G, A2614U, and G2618A. **(B)** SHAPE analysis of mini-PTC 1.4. SHAPE reactivity was plotted in the red-yellow-white spectrum where red represents high SHAPE reactivity (>1.5) and white represents low SHAPE reactivity (<0.5).

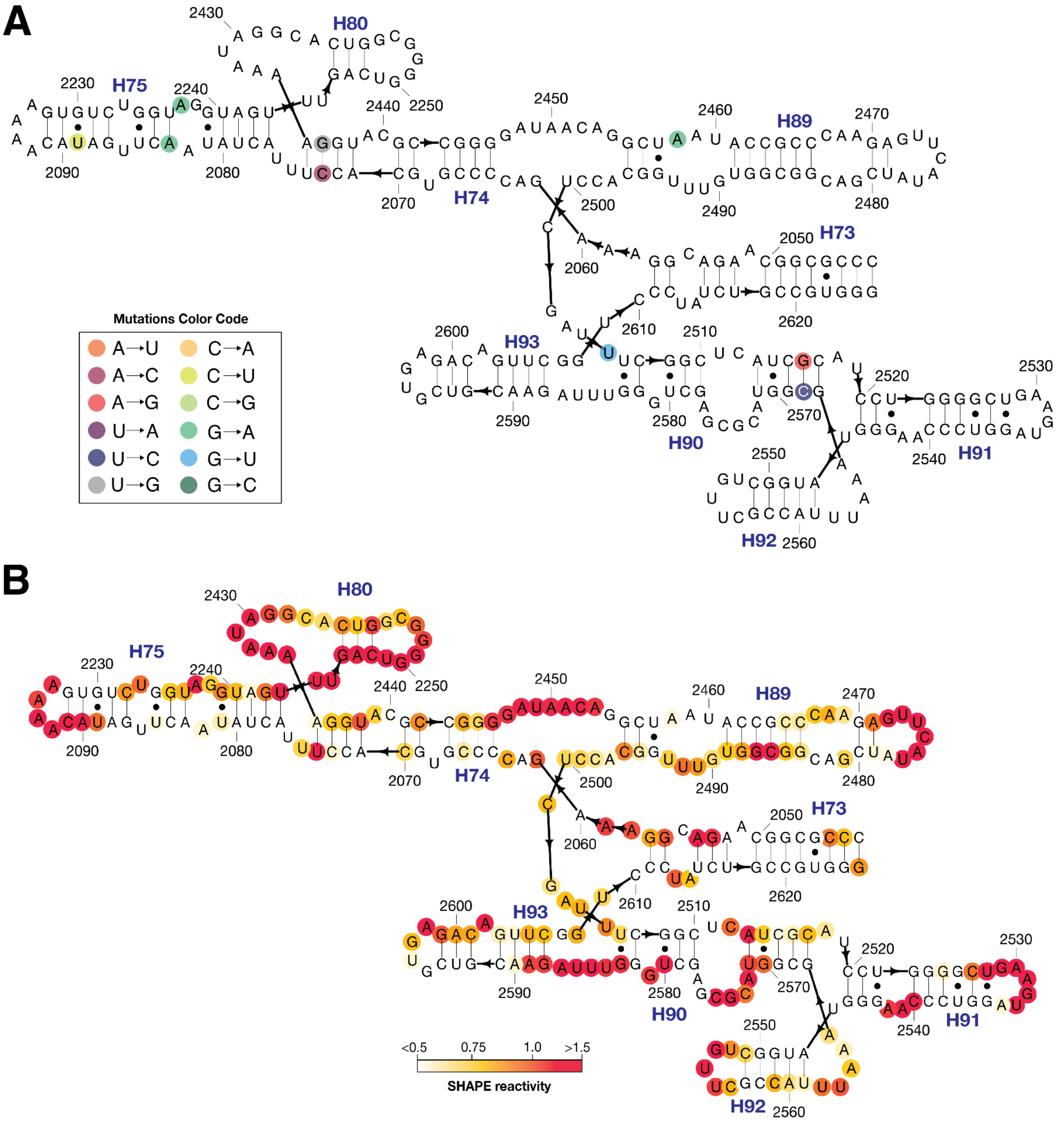


**Figure S9.** Predicted secondary structure and SHAPE analysis of mini-PTC 1.5. **(A)** Predicted secondary structure of mini-PTC 1.5. Residue numbers and helices are provided in black and blue. Mutations are highlighted in colored circles, which include mutations A2070C, G2083A, C2089U, G2237A, U2441G, G2458A, G2505U, A2516G, and U2568C. **(B)** SHAPE analysis of mini-PTC 1.5. SHAPE reactivity was plotted in the red-yellow-white spectrum where red represents high SHAPE reactivity (>1.5) and white represents low SHAPE reactivity (<0.5).

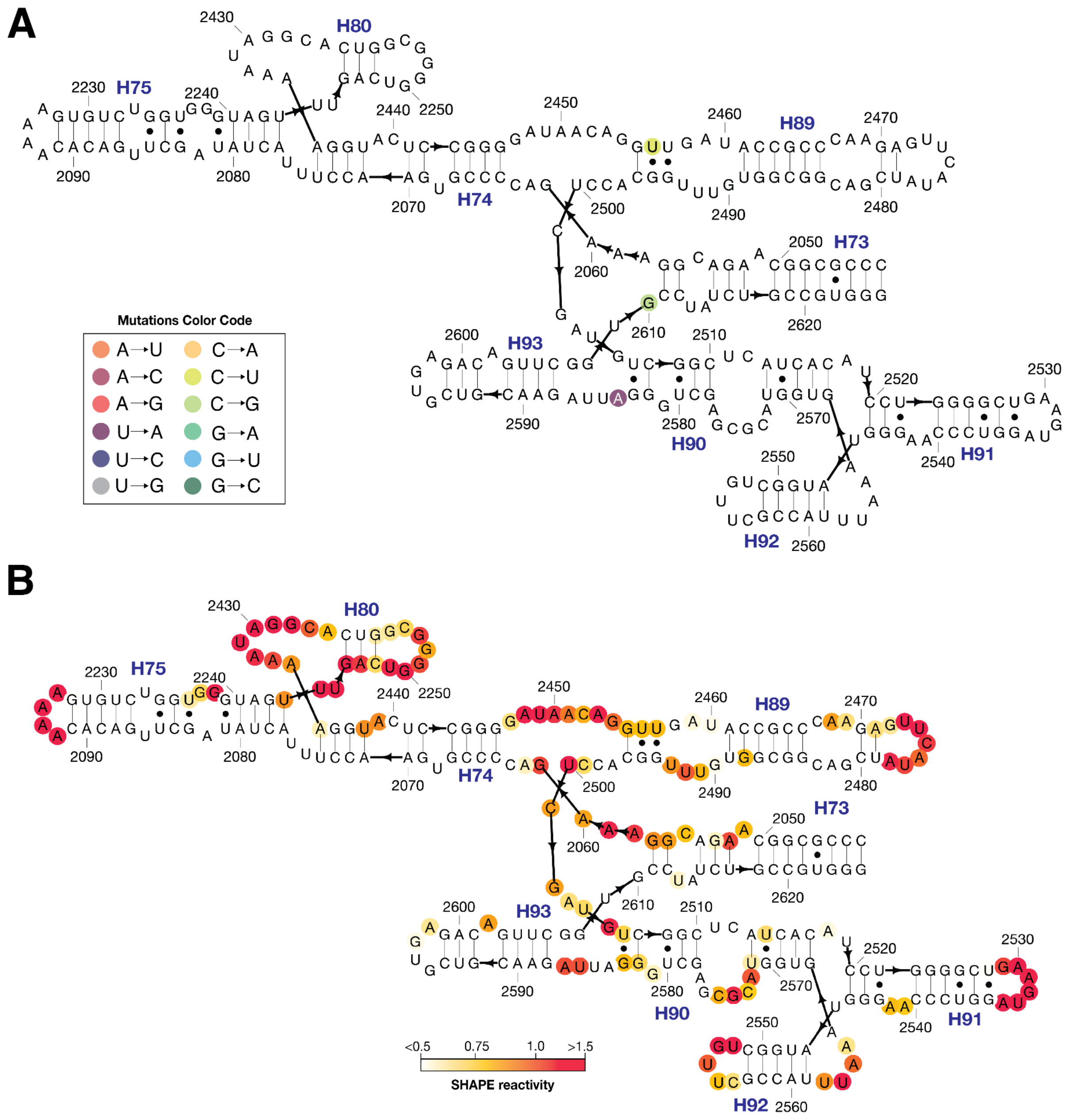


**Figure S10.** Predicted secondary structure and SHAPE analysis of mini-PTC 1.6. **(A)** Predicted secondary structure of mini-PTC 1.6. Residue numbers and helices are provided in black and blue. Mutations are highlighted in colored circles, which include mutations C2456U, U2584A, and C2610G. **(B)** SHAPE analysis of mini-PTC 1.6. SHAPE reactivity was plotted in the red-yellow-white spectrum where red represents high SHAPE reactivity (>1.5) and white represents low SHAPE reactivity (<0.5).

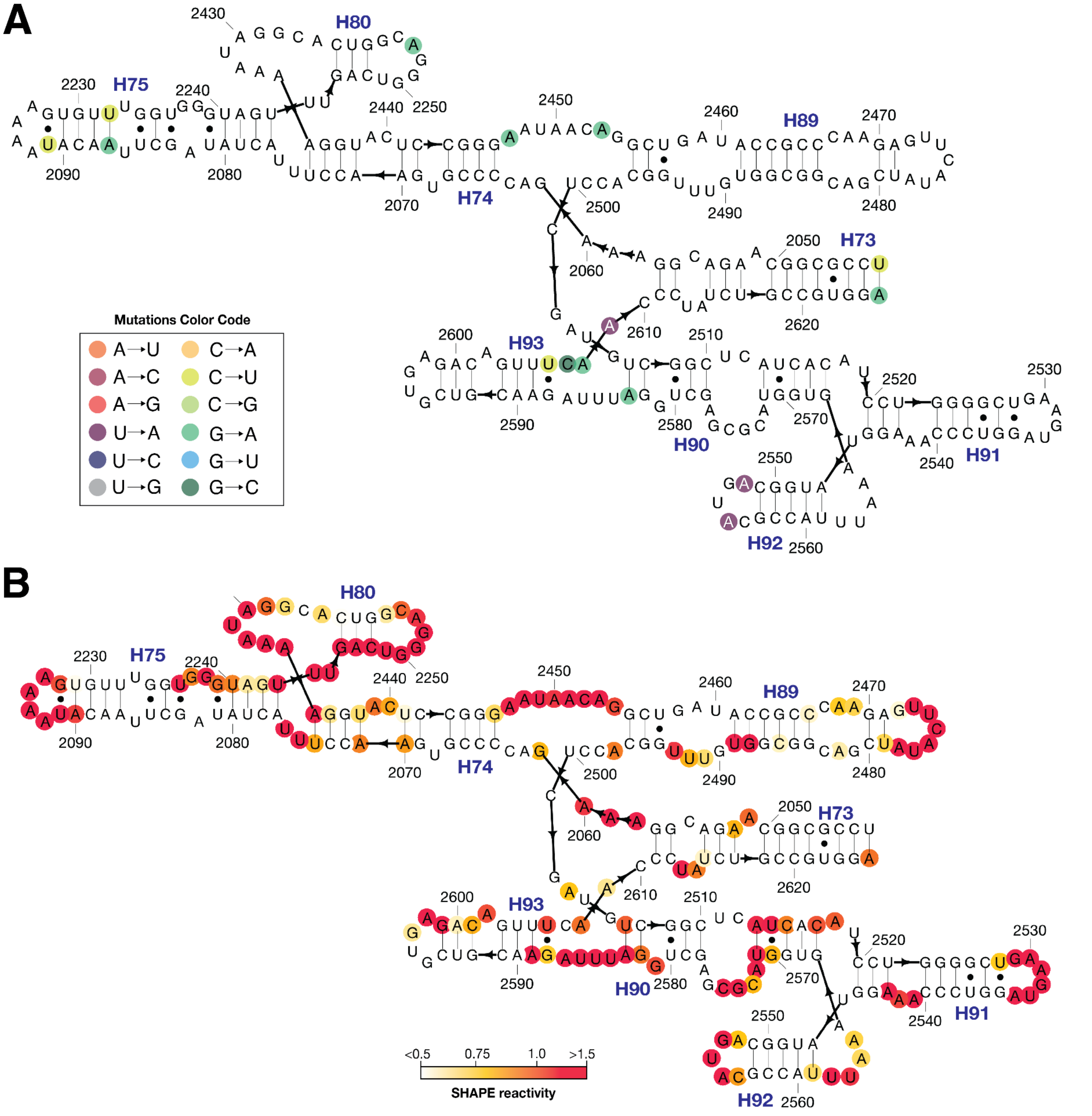


**Figure S11.** Predicted secondary structure and SHAPE analysis of mini-PTC 1.7. **(A)** Predicted secondary structure of mini-PTC 1.7. Residue numbers and helices are provided in black and blue. Mutations are highlighted in colored circles, which include mutations C2043U, G2087A, C2091U, C2232U, G2253A, G2447A, G2543A, U2552A, U2555A, G2583A, C2606U, G2607C, G2608A, U2609A, and G2625A. **(B)** SHAPE analysis of mini-PTC 1.7. SHAPE reactivity was plotted in the red-yellow-white spectrum where red represents high SHAPE reactivity (>1.5) and white represents low SHAPE reactivity (<0.5).

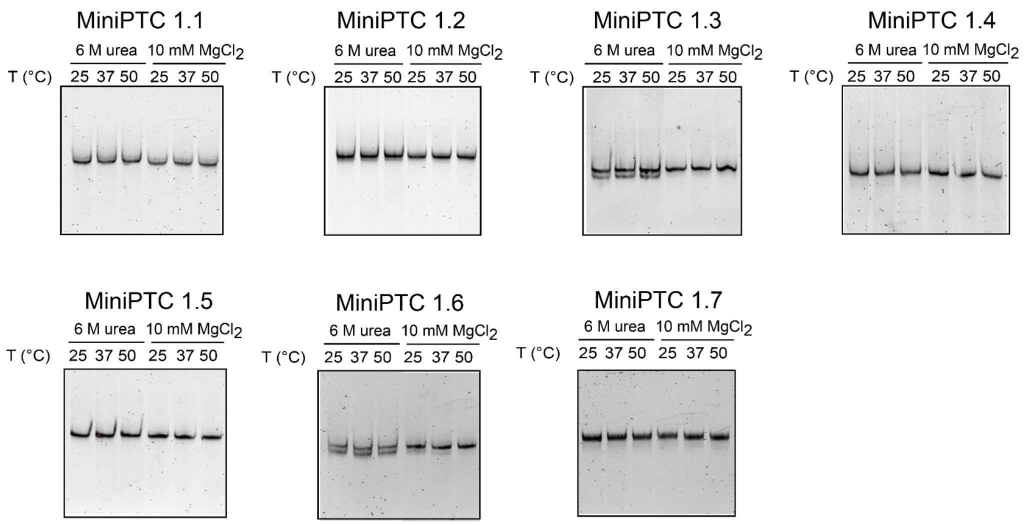


**Figure S12.** Native gel electrophoretic analysis of EteRNA-generated designs mini-PTC **1.1-1.7**. Purified samples of mini-PTC 1.1-1.7 were dissolved in 50 mM Na-HEPES, pH 8.0 supplemented with 6 M urea (lanes 1-3) or 10 mM $MgCl_2$ (lanes 4-6) and incubated at 25, 37, or 50 ºC for 20 min and applied to a 8% native acrylamide gel. All Eterna mini-PTC 1.1-1.7 migrated as a single band when re-folded in the presence of 10 mM $MgCl_2$ at 50 ºC.

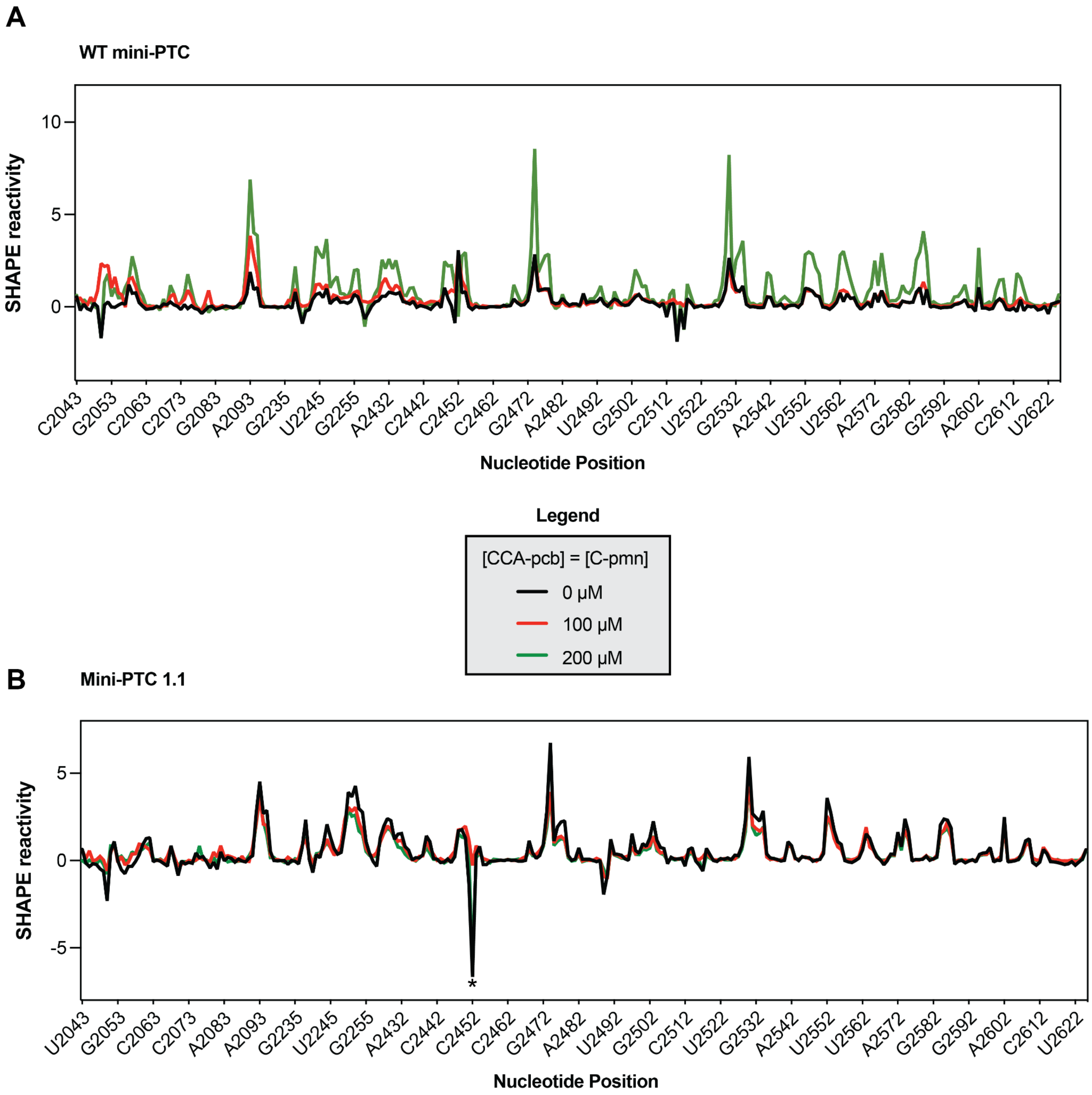


**Figure S13.** SHAPE reactivities of WT mini-PTC **(A)** and mini-PTC 1.1 **(B)** in the absence (black) or presence of 100 µM (red) or 200 µM (green) A- and P- site substrate mimics. Asterisk indicates A2451 and C2452 positions in mini-PTC 1.1 where strong negative values were observed in the absence of substrate conditions.

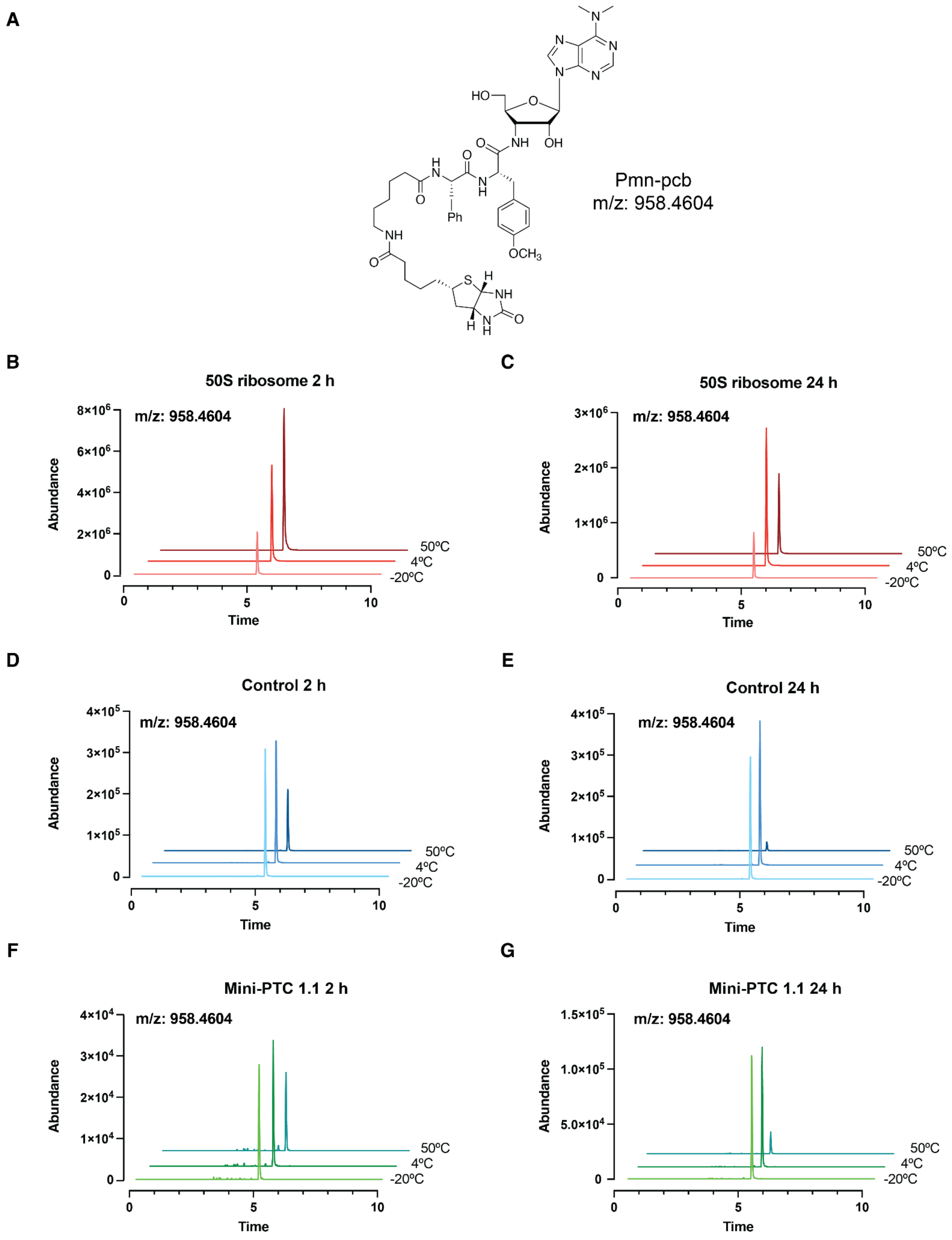


**Figure S14.** Structure and a representative of extracted ion chromatography of pmn-pcb in the fragment reactions. **(A)** Chemical structure of pmn-pcb with mass/charge ratio. **(B-C)** Extracted ion chromatography of pmn-pcb in the fragment reactions containing 50S ribosome with incubation time at 2 or 24 h. Pmn-pcb elutes at approximately t=5 min. Reactions incubated at -20 ºC, 4 ºC, and 50 ºC were represented in pink, red, and brown, respectively. **(D-E)** Extracted ion chromatography of pmn-pcb in the fragment reactions containing water with incubation time at 2 or 24 h. Reactions incubated at -20 ºC, 4 ºC, and 50 ºC were represented in light blue, blue, and dark blue, respectively. **(F-G)** Extracted ion chromatography of Pmn-pcb in the fragment reactions containing mini-PTC 1.1 with incubation time at 2 or 24 h. Reactions incubated at -20 ºC, 4 ºC, and 50 ºC were represented in light green, green, and teal, respectively.

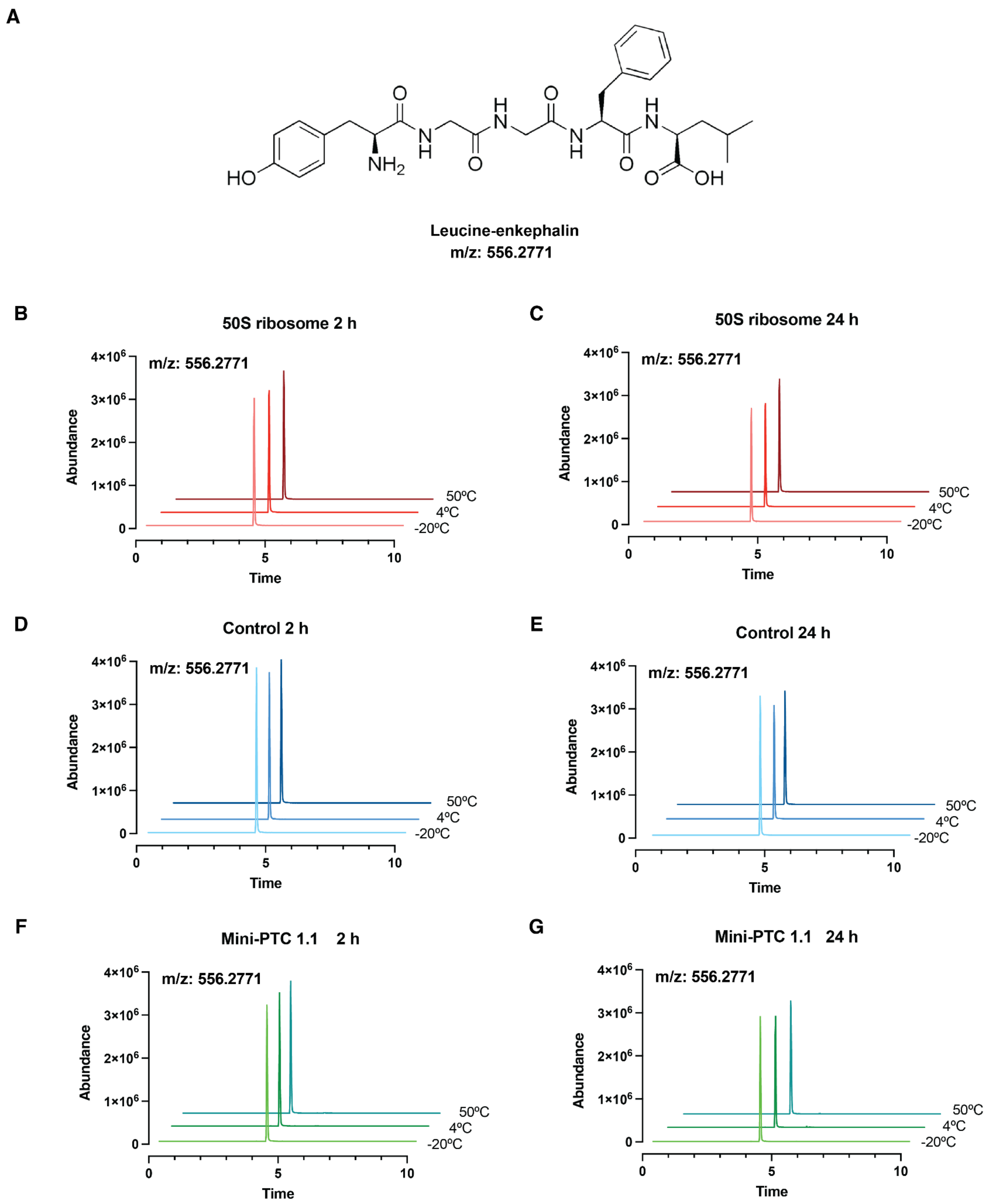


**Figure S15.** Structure and a representative of extracted ion chromatography of the co-injected leu-enkephalin (Leu-enk) in the fragment reaction mixtures, which served as an internal standard for the LC-MS quantitative analysis. **(A)** Chemical structure of Leu-enk with mass/charge ratio. **(B-C)** Extracted ion chromatography of leu-enkephalin in the fragment reactions containing *E.coli* 50S ribosomal subunit with incubation time at 2 or 24 h. Leu-enk elutes at approximately t = 4.5 min. Reactions incubated at -20 ºC, 4 ºC, and 50 ºC were represented in pink, red, and brown, respectively. **(D-E)** Extracted ion chromatography of Leu-enk in the fragment reactions containing water with incubation time at 2 or 24 h. Reactions incubated at -20 ºC, 4 ºC, and 50 ºC were represented in light blue, blue, and dark blue, respectively. **(F-G)** Extracted ion chromatography of Leu-enk in the fragment reactions containing mini-PTC 1.1 with incubation time at 2 or 24 h. Reactions incubated at -20 ºC, 4 ºC, and 50 ºC were represented in light green, green, and teal, respectively.

**Table S1** Peak area of extracted ion chromatogram of Pmn-pcb (m/z: 958.4604) in the fragment reaction at -20 ºC, 4 ºC, or 50 ºC at 2 h, and peak area of co-injected 0.04 ng leucine enkephalin internal standard (m/z: 556.2771). Activity was defined as the ratio of peak area of Pmn-pcb divided by the peak area of leucine enkephalin × 1000.

| Reaction | Time | AUC (product) rep 1 | AUC (product) rep 2 | AUC (product) rep 3 | AUC (Leu-enk) 1 | AUC (Leu-enk) 2 | AUC (Leu-enk) 3 | Ratio (product/ standard) 1 | Ratio (product/ standard) 2 | Ratio (product/ standard) 3 | Ratio x 1000 1 | Ratio x 1000 2 | Ratio x 1000 3 |
|---|---|---|---|---|---|---|---|---|---|---|---|---|---|
| Neg -20ºC | 2 h | 750056.21 | 764775.57 | 752034.9 | 9819862.85 | 9379231.08 | 9156772.27 | 0.07638153622 | 0.08153926089 | 0.08212881983 | 76.381536222 | 81.539260869 | 82.128819833 |
| Neg 4ºC | 2 h | 820410.66 | 814271.31 | 807334.2 | 8691245.28 | 8507647.84 | 8341889.36 | 0.09439506464 | 0.09571050957 | 0.09678073697 | 94.395064644 | 95.710509507 | 96.780736977 |
| Neg 50ºC | 2 h | 395014.1 | 383239.35 | 390939.95 | 8594013.54 | 8543216.78 | 8425541.5 | 0.04596386754 | 0.04485890501 | 0.04639938572 | 45.963867544 | 44.858905011 | 46.399385722 |
| PTC1.1 -20ºC | 2 h | 70315.97 | 68721.59 | 64490.36 | 8104599.89 | 8017681.79 | 7930222.42 | 0.008676056925 | 0.00857125436 | 0.00813222588 | 8.676056925 | 8.57125436 | 8.13222588 |
| PTC1.1 4ºC | 2 h | 74391.04 | 69935.27 | 66734.5 | 7931518.25 | 8034933.89 | 7952749 | 0.009379167727 | 0.00870390111 | 0.008391375108 | 9.379167727 | 8.70390111 | 8.391375108 |
| PTC1.1 50ºC | 2 h | 49724.93 | 49537.69 | 47554.6 | 7999044.61 | 7929187.76 | 7931861.3 | 0.006216358631 | 0.006247511284 | 0.005995389758 | 6.216358631 | 6.247511284 | 5.995389758 |
| 50S -20ºC | 2 h | 6400841.17 | 6353955.6 | 6315462.83 | 7876893.72 | 7732333.15 | 7765910.5 | 0.8126098177 | 0.8217384684 | 0.813228897 | 812.6098177 | 821.7384684 | 813.228897 |
| 50S 4ºC | 2 h | 15735383.09 | 15711732.36 | 15760457.86 | 7832769.52 | 7892252.28 | 7856535.56 | 2.008916904 | 1.990779286 | 2.006031506 | 2008.916904 | 1990.779286 | 2006.031506 |
| 50S 50ºC | 2 h | 24408720.71 | 24498902.9 | 24610794.19 | 8053351.67 | 8160518.57 | 8147493.81 | 3.030877293 | 3.002125746 | 3.020658225 | 3030.877293 | 3002.125746 | 3020.658225 |

**Table S2** Peak area of extracted ion chromatogram of Pmn-pcb (m/z: 958.4604) in the fragment reaction at -20 ºC, 4 ºC, or 50 ºC at 24 h, and peak area of co-injected 0.04 ng leucine enkephalin internal standard (m/z: 556.2771). Activity was defined as the ratio of peak area of Pmn-pcb divided by the peak area of leucine enkephalin × 1000.

| Reaction | Time | AUC (product) rep 1 | AUC (product) rep 2 | AUC (product) rep 3 | AUC (Leu-enk) 1 | AUC (Leu-enk) 2 | AUC (Leu-enk) 3 | Ratio (product/ standard) 1 | Ratio (product/ standard) 2 | Ratio (product/ standard) 3 | Ratio x 1000 1 | Ratio x 1000 2 | Ratio x 1000 3 |
|---|---|---|---|---|---|---|---|---|---|---|---|---|---|
| Neg -20°C | 24 h | 801752.53 | 824335.12 | 804272.45 | 8445820.15 | 8132255.27 | 7907912.82 | 0.09492891345 | 0.1013661146 | 0.1017047694 | 94.92891345 | 101.3661146 | 101.7047694 |
| Neg 4°C | 24 h | 917749 | 918502.71 | 917061.8 | 6664435.7 | 6691361.65 | 6541994.63 | 0.1377084334 | 0.1372669358 | 0.1401807632 | 137.7084334 | 137.2669358 | 140.1807632 |
| Neg 50°C | 24 h | 60516.01 | 56597.72 | 51917.89 | 7014953.19 | 7080670.1 | 7106444.88 | 0.008626716153 | 0.00799327171 | 0.00730574723 | 8.626716153 | 7.99327171 | 7.30574723 |
| PTC1.1 -20°C | 24 h | 300048.67 | 285599.01 | 288888.82 | 7026491.71 | 6964938.4 | 6767692.86 | 0.04270248687 | 0.04100524565 | 0.04268645548 | 42.70248687 | 41.00524565 | 42.68645548 |
| PTC1.1 4°C | 24 h | 289270.1 | 283597.5 | 285027.58 | 6698914.1 | 6618306.09 | 6673649.1 | 0.04318164044 | 0.04285046599 | 0.04270940466 | 43.18164044 | 42.85046599 | 42.70940466 |
| PTC1.1 50°C | 24 h | 50896.19 | 49169.8 | 50560.16 | 7039975.18 | 6939322.22 | 6861078.43 | 0.007229597932 | 0.007085677598 | 0.007369127247 | 7.229597932 | 7.085677598 | 7.369127247 |
| 50S -20°C | 24 h | 2296655.82 | 2271768.48 | 2246456.64 | 6799127.17 | 6836924.43 | 6722791.63 | 0.3377868604 | 0.3322793024 | 0.3341553277 | 337.7868604 | 332.2793024 | 334.1553277 |
| 50S 4°C | 24 h | 7871537.18 | 7838408.11 | 7789072.13 | 6537673.58 | 6582568.73 | 6566112.05 | 1.204027256 | 1.19078257 | 1.186253306 | 1204.027256 | 1190.78257 | 1186.253306 |
| 50S 50°C | 24 h | 4290304.63 | 4250627.66 | 4216983.2 | 7221582.26 | 7160914.98 | 7256112.51 | 0.5940948224 | 0.5935872262 | 0.5811628739 | 594.0948224 | 593.5872262 | 581.1628739 |
